# Decadal signatures of seasonal and ENSO-driven selection in microbial populations in distant oceans

**DOI:** 10.64898/2026.09.14.751425

**Authors:** Sergio González-Motos, Lidia Montiel, Alyse Larkin, Francisco Latorre, Caterina R. Giner, Ting-Chun Huang, Vanessa Balagué, Josep M. Gasol, Adam C. Martiny, Ramiro Logares

## Abstract

Despite the crucial role of the ocean microbiome for global ecosystem processes, its response to global change remains poorly understood. Global change will exert selection on microbial species through changes in the genetic composition of their populations, favouring strains that are better adapted to new conditions. Analyzing long-term genomic variation in microbial populations inhabiting climatically comparable but oceanographically distinct regions may provide insights into their responses to future environmental conditions. We analyzed coastal marine microbiomes from two distant long-term observatories with Mediterranean-type climates but contrasting oceanographic regimes: the Blanes Bay Microbial Observatory (BBMO; Northwestern Mediterranean Sea) and the Microbes in the Coastal Region of Orange County (MiCRO; California coast, Pacific Ocean). Sampling conducted at least monthly during 15 and 10 years, respectively, yielded 1,535 Metagenome-Assembled Genomes (MAGs) in BBMO and 1,068 MAGs plus 187 Single-Amplified Genomes (SAGs) in MiCRO. Among these, we found 250 genomes with intraspecific (>95% genome similarity) representatives occurring at both sites. In contrast, at the strain-level threshold (≥99% genome similarity), only 13 BBMO genomes matched 15 closely related representatives in MiCRO. As genome similarity increased, we observed a shift from cosmopolitan to more coastal distributions and a slight increase in genome size, pointing to niche adaptation. Across both time series, we observed widespread seasonal population structure, with most genomes (∼70%) showing significant seasonal structuring of variant composition. Moreover, in MiCRO, ∼76% of the tested genomes also showed El Niño Southern Oscillation (ENSO)-associated structure in variant composition beyond seasonal effects. Focusing on a *Prochlorococcus* genome with intraspecific representatives at both locations (>96% genome similarity) we further found recurrent seasonal mutations and non-synonymous to synonymous mutation ratios (pN/pS) from multiple genes, together with signatures of positive selection (higher pN/pS) coinciding with El Niño events in MiCRO. This suggests an imprint of ENSO in marine microbial populations. Thus, beyond seasonality, long-term climatic oscillations may shape microbial populations. This has implications for understanding how the ocean microbiome will respond to long-term environmental change and prolonged disturbances through population-level genomic variation.

## INTRODUCTION

Marine microorganisms form the foundation of ocean ecosystems and play a fundamental role in Earth’s biogeochemical cycles (*1*, *2*). They perform pivotal roles in carbon fixation, nutrient recycling, and organic matter degradation, influencing global climate and ecosystem productivity. In particular, microbes in the sunlit surface ocean are responsible for a significant portion of atmospheric carbon and nitrogen fixation (*3*, *4*). Comprising a vast diversity of bacteria, archaea, viruses, and eukaryotic microorganisms, the ocean microbiome accounts for nearly half of Earth’s primary production (*1*). Among these groups, bacteria are particularly abundant and functionally diverse, making them key players as mediators of ecosystem processes across spatial and temporal scales (*5*).

While microbial communities are often studied at broad taxonomic scales or functional groups, many of the ecological and functional processes they mediate are linked to finer scales of diversity, particularly at the population level (*6*). Populations are groups of genetically similar individuals from the same species that inhabit a specific environment at a given time (*7*, *8*). These genetic populations, sometimes referred to as “genetic clusters,” are distinguishable based on their cohesive genomic features and ecology. Within a bacterial species, multiple populations can occur, each defined by distinct traits and genomic features (*9*). As such, populations represent fundamental ecological and evolutionary units upon which natural selection acts. Because of this, microbial populations may respond rapidly to environmental changes, making them sensitive sentinels of ecosystem shifts (*10*). Decoding their structure, dynamics, and genomic variation is essential to understanding how microbial life responds to environmental change (*6*). Despite their ecological relevance, our understanding of how oceanic microbial populations respond to long-term environmental change remains limited. Most knowledge of marine microbiomes stems from snapshot surveys (*11*, *12*) or long-term studies that capture taxonomic composition and functional traits (*13–15*). However, long-term patterns in microbial population structure, genomic variation, and evolutionary responses are less well characterized. In particular, the ecological and evolutionary processes (mutation, selection, gene flow, and drift) that structure marine microbial populations over multiple years or decades remain largely underexplored (*16*, *17*).

In temperate and high-latitude ecosystems, seasonal fluctuations in temperature, light, and nutrient availability are well known to structure microbial communities across a range of ocean environments (*18–21*). Multiple studies have shown that microbial taxonomic composition and functional traits vary predictably over the course of the year, with recurring seasonal patterns observed in coastal and open-ocean systems alike (*14*, *15*, *22*). For example, the relative abundance of phototrophic, heterotrophic, and mixotrophic taxa often shifts seasonally in response to changes in light regime, stratification, and nutrient availability (*23*). Although usually described at broader taxonomic scales, these patterns ultimately arise from population-level processes, such as changes in the abundance or genomic composition of individual lineages, which remain less well understood. In particular, we know only partially of the seasonal selection pressures acting on microbial populations and their evolutionary consequences. Seasonal selection pressures drive recurrent shifts in community and genomic composition, including changes in allele frequencies and ecotype configurations (*24–26*). For example, seasonal abundance patterns have been documented among ecotypes of SAR11, *Synechococcus*, and *Prochlorococcus*, where distinct lineages dominate under different temperature or nutrient regimes (*27*, *28*). Also in the nitrogen-fixing symbiont UCYN-A, whose ecotypes display distinct seasonal niches linked to their haptophyte hosts (*29*) or among marine *Thaumarchaeota*, whose ecotypes track seasonal changes in water column structure and nutrient conditions (*30*, *31*). Altogether, these examples highlight the fine-tuning of microbial population diversity to seasonal environmental heterogeneity.

Beyond seasonal cycles, ocean microbes may also experience other types of large-scale environmental variations, such as the El Niño-Southern Oscillation (ENSO) in the Pacific Ocean, which generates anomalous warming and nutrient shifts that propagate across marine systems (*32*). El Niño begins when the usual east-to-west trade winds over the equatorial Pacific weaken or reverse, allowing bursts of westerly winds to drive warm waters from the western Pacific eastward (*33*, *34*). By modifying environmental conditions, El Niño can affect the composition of microbial communities. For instance, the 2015 El Niño, one of the strongest on record, produced exceptional warming across the Eastern North Pacific (*35*), leading to increased cyanobacterial abundances and major changes in zooplankton community structure within the California Current (*36*, *37*). Recent work has shown that El Niño induced a marked shift in both the taxonomic and functional structure of the microbiome, including increased prevalence of oligotrophic cyanobacteria, reduced abundance of genes involved in organic matter degradation, and elevated signals of nutrient stress, suggesting a transition to a summer-like, stratified state (*38*). In addition to these community-level changes, it has also been documented that common microbial taxa such as SAR11, *Synechococcus*, and *Prochlorococcus* exhibited shifts in ecotype composition, transitioning from cold-to warm-adapted lineages during El Niño conditions (*39*, *40*). These studies, which rely on marker gene data and reference genome recruitment, provide compelling evidence of lineage-level responses, showing that both seasonal cycles and ENSO-driven anomalies can structure microbial communities and influence population-level dynamics. However, they do not resolve fine-scale genetic changes within the populations, and it remains unclear how the interplay of these environmental fluctuations drives the structure of microbial populations across regions. Addressing this question requires comparing microbial populations that are found across oceanographically distinct regions.

Analyzing variants of the same microbial genome across distinct marine regions enables a direct comparison of how these microbes respond to different climatic or ecological regimes. Such comparisons are rare and offer unique opportunities to investigate fine-grained adaptation and the genomic regions under selection. A threshold of 95% Average Nucleotide Identity (ANI) is commonly used to define species-level similarity, while 99% ANI is used to delineate strains (*41*, *42*). These ANI-based clusters are widely used to define *metagenomic populations*, the operational genomic units for studying microbial adaptation in natural communities (*43*), and provide a practical framework to investigate evolutionary dynamics across sites. Previous studies have shown that specific prokaryotic taxa, such as *Prochlorococcus* and SAR11, are globally distributed at the species level (*11*, *44*, *45*), but this does not indicate whether the same strains persist across sites. Indeed, single-cell genomics has revealed that even within a single marine location, *Prochlorococcus* harbors hundreds of coexisting subpopulations, a microdiversity that contrasts sharply with the apparent global homogeneity inferred from species-level patterns (*46*). In prokaryotes, growing evidence indicates that strain-level diversity often has clear biogeographic structure, meaning that even when species appear globally distributed, their strains differ across locations. For example, globally distributed taxa such as SAR11 harbor distinct regional strain clusters, with fine-scale genomic differentiation across oceanic provinces that appear to be promoted by adaptation despite broad species-level ubiquity (*47*). Furthermore, biogeographic structuring below broad taxonomic levels has also been reported in other marine planktonic groups: in marine DNA viruses, intra-population genetic variation and community composition differ across major ecological zones of the global ocean, consistent with geographically structured microdiversity (*48*); and in eukaryotic plankton, genome-resolved analyses across surface oceans have likewise revealed broad biogeographic organization and recurrent functional convergence among widespread lineages (*49*). However, no studies to date have used long-term time-series data from geographically distant ecosystems to track microbial populations present in both locations and through time. The recovery of identical or near-identical prokaryotic genomes across large spatial and temporal scales offers an opportunity to investigate how seasonality and long-term environmental variability, such as ENSO, may shape microbial populations.

Here, we leverage two long-term coastal marine time series to investigate microbial population dynamics across contrasting oceanographic regimes. The first time series originates from the LTER Blanes Bay Microbial Observatory (BBMO) in the northwest Mediterranean Sea, off the coast of Catalonia, Spain, where coastal microbial communities have been sampled monthly for over 15 years. BBMO represents a markedly seasonal microbiome from a coastal oligotrophic site (*50*). The second time series is the ‘Microbes in the Coastal Region of Orange County’ (MiCRO), a decade-long observatory on the eastern coast of the Pacific Ocean in California, USA. Both observatories are located in coastal regions with Mediterranean-type climates. However, MiCRO lies within the Southern California Bight (SCB), a critical transition zone between subpolar and subtropical biomes where the effects of large-scale climatic oscillations, such as the El Niño-Southern Oscillation (ENSO), are particularly pronounced. The SCB is shaped by the interplay of two contrasting ocean currents: the southward-flowing California Current, which brings cold, low-salinity, and nutrient-poor waters, and the northward-flowing California Undercurrent, which carries warmer, saltier, and nutrient-rich waters (*51*). Like BBMO, MiCRO displays strong seasonal patterns, yet is further affected by high interannual variability primarily driven by ENSO cycles.

Using high-resolution metagenomic data from short- and long-read sequencing technologies as well as Single-Cell Genomics, we reconstructed thousands of microbial genomes, including Metagenome-Assembled Genomes (MAGs) and Single-Amplified Genomes (SAGs), enabling comparative population genomics across sites. We focused on identifying genomes and strains present in both of these distant regions, characterizing their seasonal and long-term dynamics, and examining evolutionary patterns through variant analysis, in particular the pN/pS ratio. This ratio compares the frequency of non-synonymous (amino acid-changing) to that of synonymous (silent) mutations in genes and serves as a proxy for the strength and direction of natural selection (*52*). In particular, we tested whether large-scale climatic oscillations, such as the El Niño Southern Oscillation (ENSO), shape population dynamics when comparing species or strains present in both time series. This comparative framework offers new insights into the stability, adaptability, and climatic sensitivity of ocean microbes over decadal scales.

## RESULTS

### Environmental regimes and climatic variability across the two time-series

The two time-series analyzed in this study are located in distinct coastal marine environments: the Blanes Bay Microbial Observatory (BBMO), in the northwestern Mediterranean Sea (41°40′N, 2°48′E), and the Microbes in the Coastal Region of Orange County (MiCRO), in Newport Beach, Pacific Ocean (33.608°N, 117.928°W) at similar latitudes (**Figure 1A**). Both sites are characterized by strong seasonal variability in surface seawater temperature. At BBMO, temperatures ranged from ∼12°C in winter to ∼25°C in summer, following a regular annual cycle over the 15-year sampling period (**Figure 1B)**. Similarly, at MiCRO, seasonal fluctuations were observed, with temperatures ranging between ∼13°C and ∼22°C each year (**Figure 1C**).

**Figure 1.**
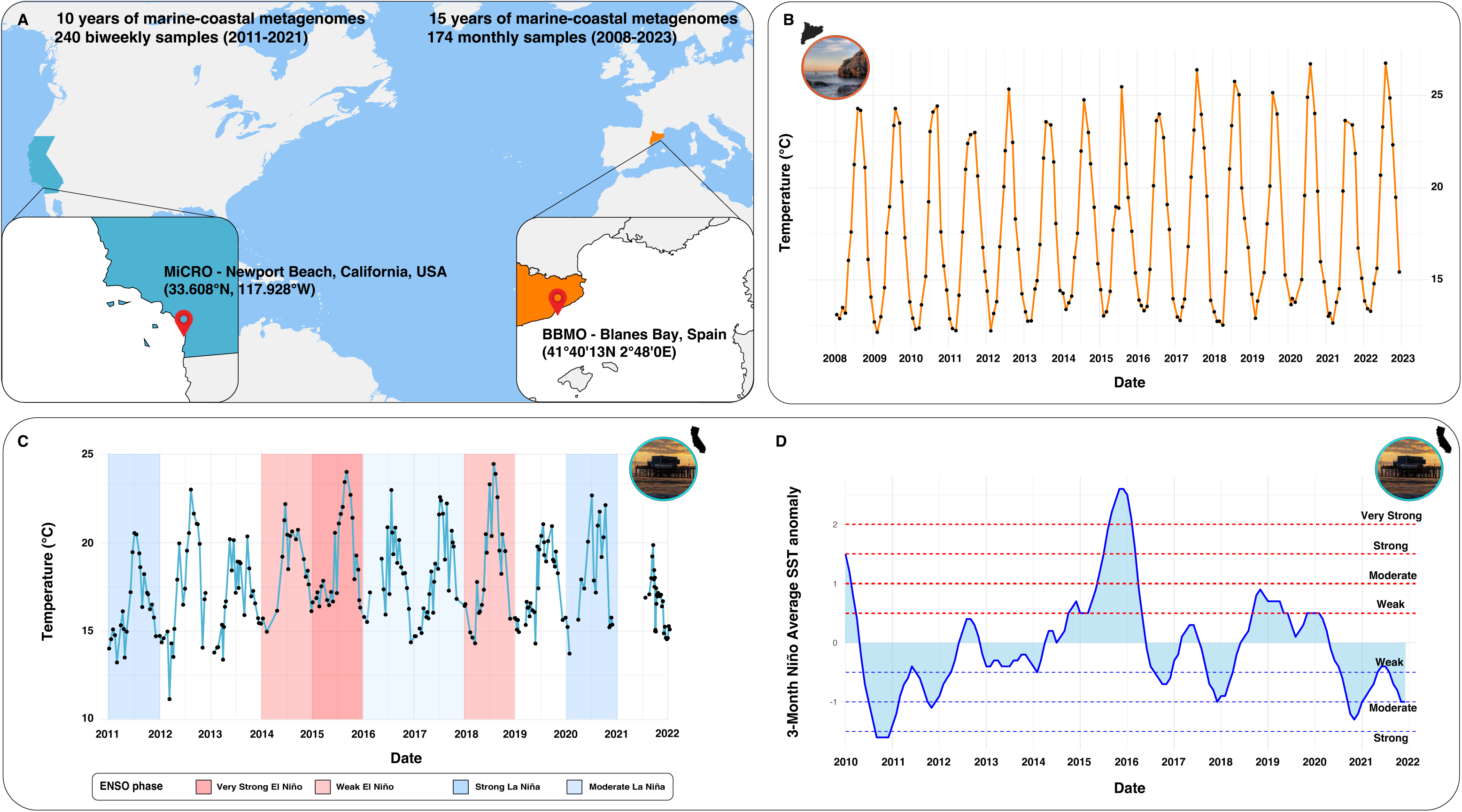
Seasonal temperature patterns and climatic variability across BBMO and MiCRO. **Panel A.** Map showing in orange the location of the Blanes Bay Microbial Observatory (BBMO), in the northwest Mediterranean Sea, Catalonia, Spain (41°40′N, 2°48′E) and in blue the Microbes in the Coastal Region of Orange County (MiCRO), in Newport Beach, California, USA (33.608°N, 117.928°W). **Panel B.** Seasonal Sea Surface Temperature (SST) at BBMO over 15 years, illustrating the regular annual cycle from ∼12°C in winter to 24-25°C in summer. **Panel C.** Seasonal SST at MiCRO over 10 years, ranging between ∼12°C and ∼22°C annually, with anomalous warming up to ∼25°C during El Niño years. Note that ENSO phases according to NOAA are indicated in colours in the corresponding MiCRO years. **Panel D.** Oceanic Niño Index (ONI) through time based on NOAA Niño 3.4 SST anomalies, highlighting very strong El Niño events in 2015-2016 (ONI > +2°C).

In addition, MiCRO exhibited clear interannual variability associated with the El Niño Southern Oscillation (ENSO), as expected for a microbiome located within the Pacific Ocean temperate ocean. The Oceanic Niño Index (ONI index), based on the 3-month averages of sea surface temperature (SST) anomalies in the Niño 3.4 region (5°N-5°S, 120°-170°W), revealed strong El Niño events in 2015-2016, with ONI values exceeding +1.5°C according to NOAA data (*53*) (**Figure 1D**). These events coincided with temperature anomalies at MiCRO, where summer maxima reached ∼24°C during El Niño years, exceeding the typical seasonal maximum of ∼22°C. In contrast, BBMO displayed relatively stable interannual temperature dynamics, with no pronounced warming anomalies across the time series. Together, these records indicate that while both BBMO and MiCRO are seasonally structured systems, MiCRO experiences additional interannual forcing linked to large-scale climatic oscillations. This environmental contrast provides a valuable backdrop for comparing microbial population structure and dynamics under different temporal modes of environmental variability.

### Species and strains present in both time series

We recovered a total of 1,535 Metagenome-Assembled Genomes (MAGs) from the Blanes Bay Microbial Observatory (BBMO) samples and 1,068 MAGs together with 187 Single-Amplified Genomes (SAGs) from Newport (MiCRO) samples, using both short- and long-read sequencing data. To determine whether similar genomes were present in these two geographically distant locations, we performed dereplication of all genomes at two different Average Nucleotide Identity (ANI) thresholds: 95% (species-level) and 99% (strain-level) (*41*, *42*).

At 95% ANI, the dereplication analysis initially revealed 615 genomes grouped into clusters shared between BBMO and MiCRO (**Figure 2A**). Because high ANI values can result from partial alignments, we further applied minimum pairwise alignment coverage thresholds. This retained 458 shared genomes at ≥50% coverage, 335 at ≥60% coverage, and 250 at ≥70% coverage. Hereafter, we refer to these as shared genomes, defined as genomes recovered independently from the two sites that grouped into the same dereplicated cluster at the specified ANI and alignment coverage thresholds. Unless otherwise stated, downstream analyses were based on the conservative set retained at ≥95% ANI and ≥70% alignment coverage (120 BBMO genomes and 130 MiCRO genomes). These genomes represent putative species that are present in both coastal environments despite the large geographical separation. Among these 250 shared genomes, 120 were assembled from BBMO and 130 from MiCRO, indicating that multiple genomes from one observatory could dereplicate into a single one from the other. The remaining genomes were unique to each site: 1415 were exclusive to BBMO, and 1125 were exclusive to MiCRO. This pattern suggests a distinct local genomic pool in each region, with a substantial set of species shared between them. The slightly larger contribution from MiCRO to the shared pool may reflect the presence of more cosmopolitan lineages in MiCRO compared with BBMO. Differences in sampling depth and genome recovery may also contribute to the observed asymmetry in shared genomes.

**Figure 2.**
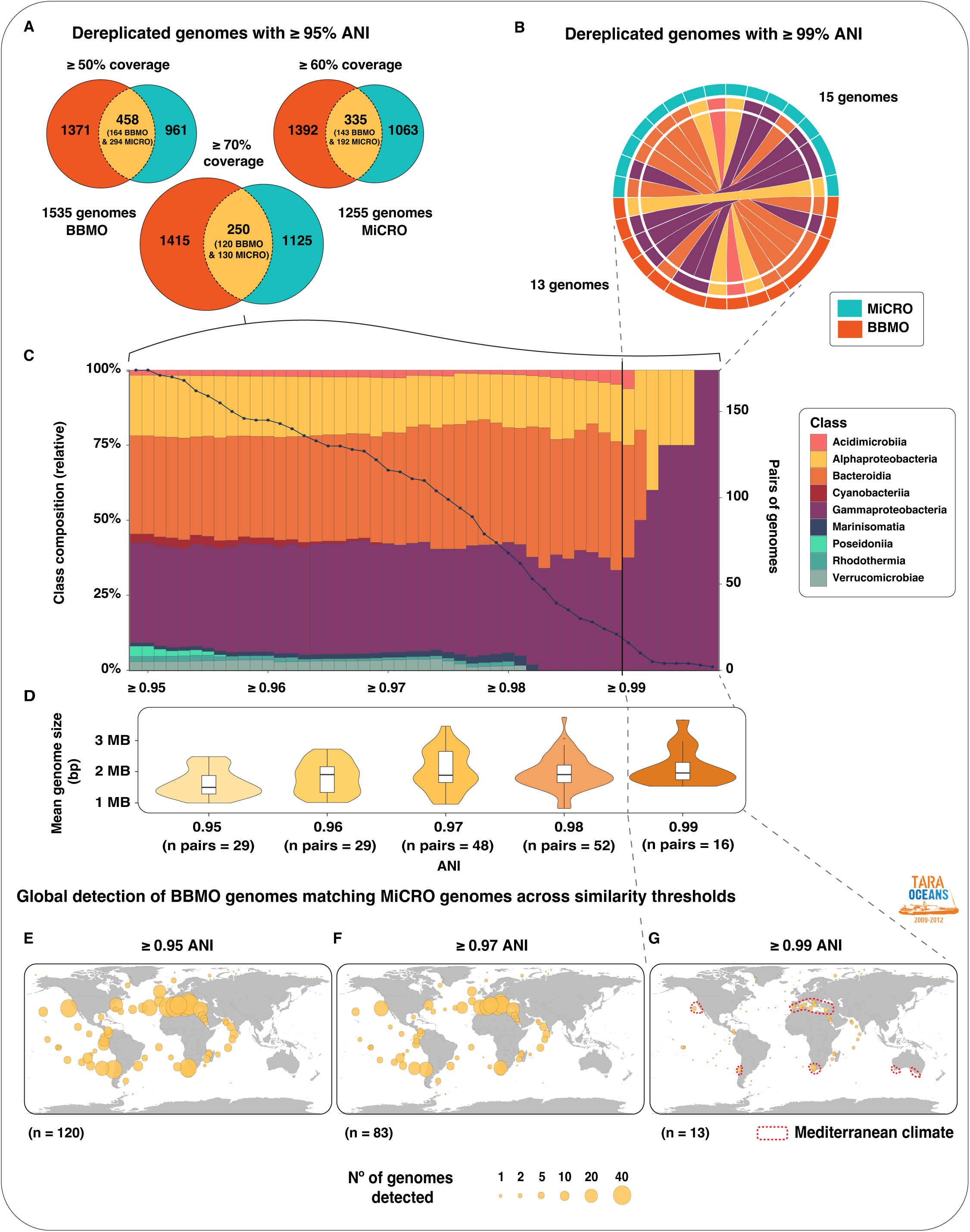
Considerable species-level overlap but pronounced strain-level divergence between BBMO and MiCRO, with similarity-dependent taxonomic and biogeographic structuring between locations. **Panel A.** Species-level overlap of genomes (yellow) recovered from Blanes Bay Microbial Observatory (BBMO) in orange and MiCRO (Newport, California) in blue after dereplication at 95% Average Nucleotide Identity (ANI). Venn diagrams show shared genomes retained at ≥50%, ≥60%, and ≥70% alignment coverage. The conservative ≥70% coverage threshold retained 250 shared genomes, including 120 BBMO and 130 MiCRO genomes, and was used for downstream analyses. Genomes included both Metagenome-Assembled Genomes (MAGs) and Single-Amplified Genomes (SAGs) reconstructed from long- and short-read sequencing data. **Panel B.** Strain-level overlap at 99% ANI, revealing only 13 BBMO genomes (in orange) dereplicating with 15 MiCRO genomes (in blue). **Panel C.** Taxonomic relative composition at class level of shared genomes across ANI thresholds depicted in colours. Within each ANI threshold, all genome pairs with equal to or greater than the indicated value are shown, with the number of pairs also indicated by a cumulative line. **Panel D.** Mean genome size in base pairs (bp) of shared genomes across increasing ANI thresholds. Each violin plot represents the mean genome sizes of shared genomes grouped by successive ANI intervals. **Panels E-G.** Global detection of BBMO-MiCRO shared genomes across increasing genome similarity thresholds. Maps show the cumulative number of shared genomes detected across *Tara* Oceans metagenomes for thresholds of ≥95% ANI (E), ≥97% ANI (F), and ≥99% ANI (G), illustrating a progressive shift from broadly distributed species-level genomes to regionally constrained strain-level genomes.

When applying a more stringent threshold of 99% ANI to examine fine-scale diversity, corresponding to strain-level resolution, we identified only 13 BBMO genomes that dereplicated with 15 MiCRO genomes (**Figure 2B**). This considerable reduction indicates that despite sharing hundreds of species-level taxa, the actual strains present in each site are largely distinct. At the species level (95% ANI), shared genomes were broadly distributed across multiple bacterial lineages, including *Acidimicrobiia*, *Alphaproteobacteria*, *Bacteroidia*, *Cyanobacteria*, *Gammaproteobacteria*, *Marinisomatia*, *Poseidoniia*, *Rhodothermia* and *Verrucomicrobiae* (**Figure 2C**). In contrast, applying a strain-level threshold (99% ANI) substantially reduced this taxonomic breadth, with shared genomes restricted to a smaller subset of taxa, primarily within Alphaproteobacteria (including Puniceispirillaceae, *Lentibacter*, *Planktomarina*, and Thalassobaculales) and Gammaproteobacteria (including *Glaciecola*). Several taxonomic groups that were shared at the species level were no longer represented at the strain level, indicating a fine-scale genomic divergence across different lineages.

Genome size differed significantly across ANI thresholds (Kruskal-Wallis, p = 7.01 × 10^-3^) and showed a positive correlation with ANI, with genomes shared at lower ANI thresholds (95% ANI) tending to be smaller than those shared at higher thresholds (99% ANI) (Pearson r = 0.253, p = 7.527 x10^-4^, **Figure 2D**). This pattern suggests that shared genomes in the lower part of the similarity spectrum may be enriched in lineages with streamlined genomes, which are typical of the open oligotrophic ocean (*54*). In turn, shared genomes of strains in the upper similarity range may be enriched in taxa with expanded gene repertoires, potentially associated with meso or eutrophic habitats, such as coastal areas like BBMO and MiCRO (*55*). To further examine whether genomes associated to different lifestyles (oligotrophs vs. meso/eutrophic taxa) are present across the ANI similarity range of shared genomes, we investigated their global distributions across ANI thresholds using *Tara* Oceans metagenomes (**Figure 2E-G**). We hypothesized that shared genomes in the lower similarity range may include a higher representation of oligotrophs, with small genomes and broad geographic distributions, while shared genomes in the upper similarity range (i.e., shared strains) could be enriched in larger genomes that could be more typical of coastal areas (that is, geographically more restricted) that tend to have a higher concentration of nutrients than the open ocean. At the species level (lower range of ANI; ≥95%), shared genomes were widely detected across the global ocean, with high occurrence across multiple oceanic basins and both coastal and open-ocean regions (**Figure 2E**). Increasing the ANI threshold to ≥97% reduced the number of genomes detected in each sample but the same broad pattern remained (**Figure 2F**). At the strain level (≥99% ANI), shared genomes exhibited a markedly restricted biogeographic pattern (**Figure 2G**). These genomes were predominantly detected in coastal regions, with a strong enrichment in Mediterranean-climate areas, and were largely absent from oligotrophic open-ocean regions. These results indicate that increasing genomic similarity progressively shifts the biogeographic signal from globally distributed species to regionally constrained strains, which also differ in genome size.

Beyond identifying shared species and strains between BBMO and MiCRO, we examined how their abundances vary over time at each site. Among the MAGs and SAGs present at both locations, we selected representative genome pairs to compare their temporal dynamics. These included high-quality genomes from Thalassobaculales and Ca. *Micropelagos* (ANI ≥ 98%, completeness ≥ 94%), as well as iconic taxa such as Flavobacteriaceae, SAR86, *Synechococcus*, and *Prochlorococcus*. In general, many shared genomes showed consistent seasonal abundance patterns across BBMO and MiCRO with comparable abundance. Genomes that peaked in winter or summer in one time-series tended to do so at the other, suggesting that similar seasonal cues structure microbial communities in both ecosystems, despite their geographic separation (ca. 14,403 Km minimum distance by sea) as depicted in **Table 1**.

**Table 1.** Seasonal phenology metrics of representative genomes present in BBMO and MiCRO time series.

|  | *BBMO*<br>genome | *MiCRO*<br>genome | *Peak*<br>*BBMO* | *Peak*<br>*MiCRO* | *Phase lag*<br>(months) | *Amplitude*<br>*BBMO* | *Amplitude*<br>*MiCRO* |
| --- | --- | --- | --- | --- | --- | --- | --- |
| *Thalassobaculales* | bin.G3.66 | 3300046169_16 | Oct | Oct | 0 | 1.32 | 0.84 |
| *Micropelagos* | bin.G2.101 | 3300047475_11 | Jun | Aug | 2 | 8.50 | 1.70 |
| *Synechococcus* | bin.G4.469 | UncmicAM_433_B22_FD | Apr | Jul | 3 | 13.60 | 1.90 |
| *SAR86* | bin.G2.329 | 3300046170_199 | May | Sep | 4 | 3.06 | 2.50 |
| *Flavobacteriaceae* | bin.G2.263 | 3300047157_210 | Mar | Jan | -2 | 2.50 | 0.87 |
| *Prochlorococcus* | BL_0908_bi<br>n.full.869 | UncmicAM_433_B21_FD_<br>contigs | Nov | Sep | -2 | 154 | 5.23 |

A clear example of this is observed in the Thalassobaculales and Ca. *Micropelagos* genomes, which consistently showed higher abundance during the warmer months in both ecosystems **(Figure 3A and B)**, with nearly identical peak timing for Thalassobaculales (October at both sites) and a modest phase lag of approximately two months for Ca. *Micropelagos* **(Table 1)**. Despite this general synchrony, some genomes exhibited site-specific peaks or marked differences in abundance, pointing to local abiotic or biotic effects. For instance, the *Synechococcus* genome pair shows sharp, episodic peaks in BBMO, while maintaining a more continuous, moderate presence in MiCRO (**Figure 3C**), accompanied by a seasonal delay of approximately three months in peak timing at MiCRO (**Table 1**). Similarly, the SAR86 genome pair exhibited a pronounced phase shift, with peak abundances in warm months at BBMO but peaking roughly four months later at MiCRO (**Figure 3D**; **Table 1**). Lastly, among the shared representative genomes, two pairs, one belonging to Flavobacteriaceae and the other to *Prochlorococcus*, exhibited seasonal dynamics comparable to those described above, with consistent but site-specific timing of peak abundance (**Table 1**). Beyond these seasonal patterns, both pairs of genomes showed contrasting interannual abundance patterns in MiCRO that appeared to align with El Niño dynamics (**Figure 4**). In MiCRO, the Flavobacteriaceae genome abundance tended to decrease during years with high ONI index values (e.g., 2015-2016) and was strongly negatively associated with the Oceanic Niño Index (Pearson r = -0.46, p = 1.8 × 10^-7^). This suggests sensitivity to El Niño-related surface warming and associated changes in water-column stratification and nutrient availability. In contrast, *Prochlorococcus* abundance in MiCRO increased during or right after these same El Niño years, with pronounced peaks in 2015, 2016, and 2019 (**Figure 4B**), consistent with a positive association with the Oceanic Niño Index (Pearson r = 0.29, p = 0.001). In turn, at BBMO, both genomes followed more regular seasonal dynamics, with no significant association with the Oceanic Niño Index (**Figure 4**). Notably, *Prochlorococcus* abundance was substantially higher in BBMO overall than in MiCRO (ca. one order of magnitude; **Figure 4B**).

**Figure 3.**
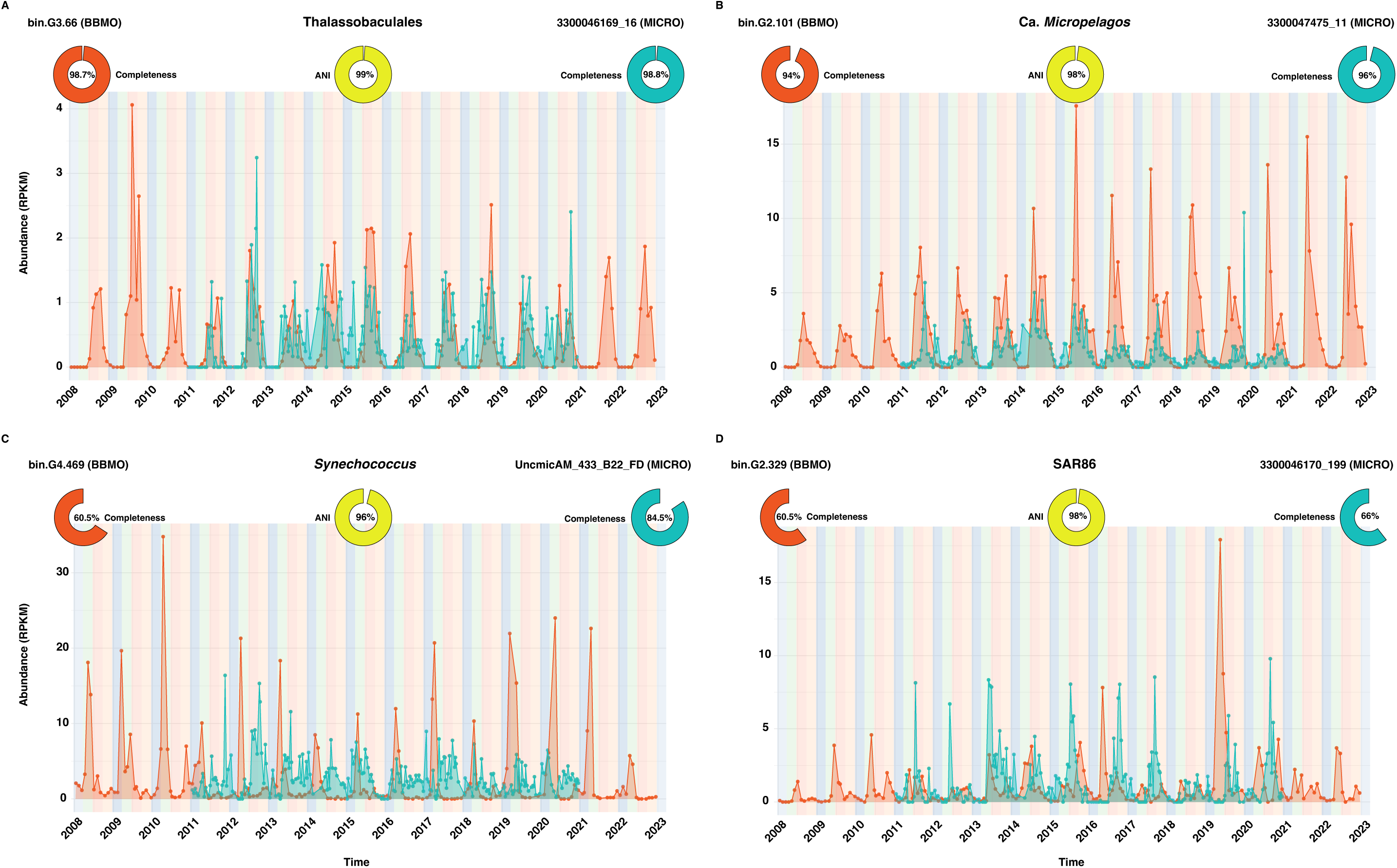
Temporal dynamics of selected genomes detected in both BBMO and MiCRO. Abundance profiles in Reads Per Kilobase per Million mapped reads (RPKM) through time for selected high-quality genome pairs present in both time series. Genome completeness (orange circles for BBMO and blue circles for MiCRO genomes) and ANI values (yellow circle) are shown in each panel. **Panel A and B.** Thalassobaculales and Ca. *Micropelagos* genomes display consistent seasonal abundance peaks in summer across both sites. **Panel C.** *Synechococcus* shows sharp and episodic peaks at BBMO but maintains a more continuous and moderate presence in MiCRO. **Panel D.** SAR86 exhibits a phase shift, with peak abundances during warmer months occurring at different times between the two sites. Together, these patterns suggest that while specific genomes present in both sites show comparable seasonal dynamics across BBMO and MiCRO, others display site-specific differences in amplitude or timing, reflecting both shared environmental cues and local effects.

**Figure 4.**
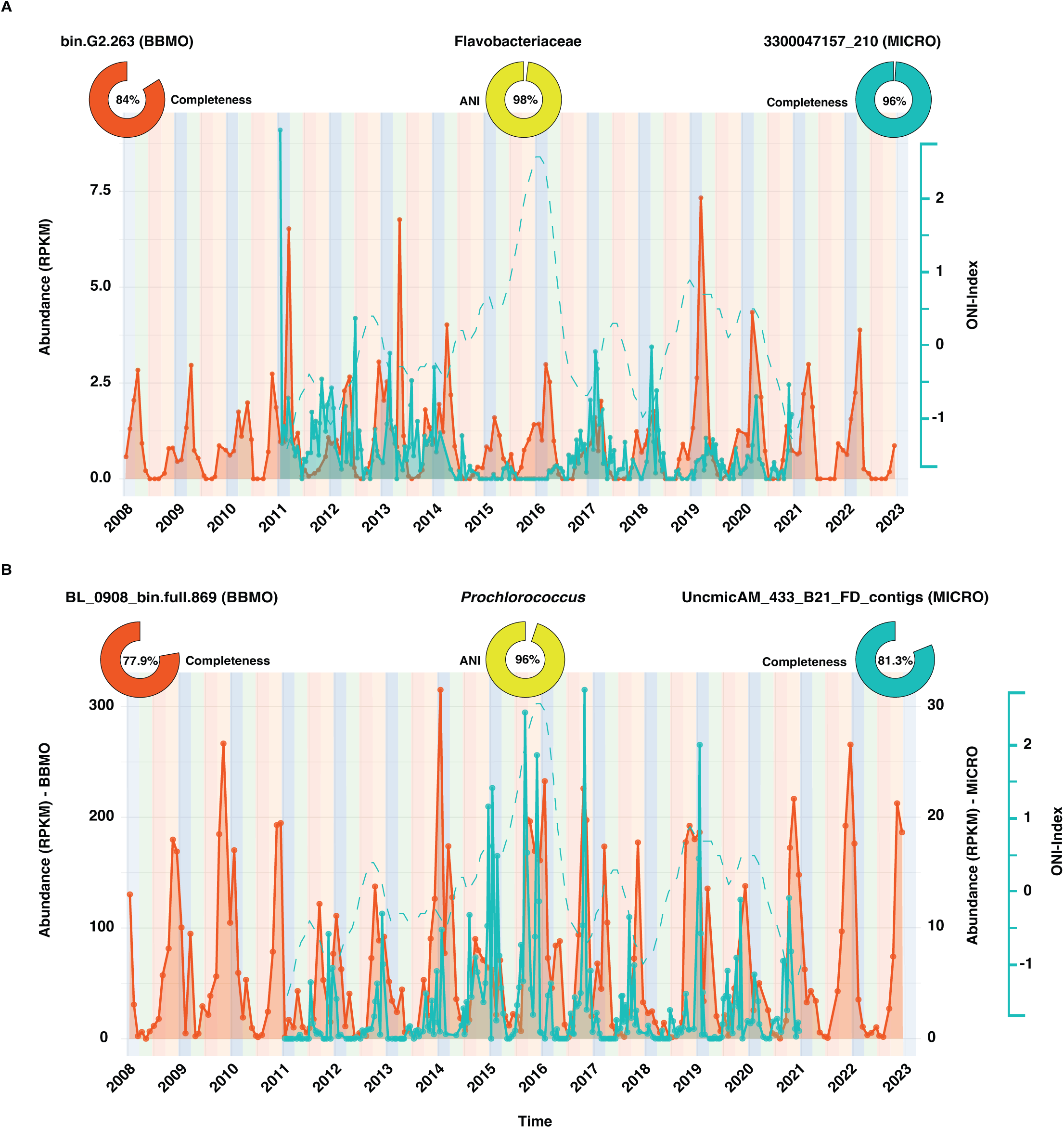
Temporal dynamics of Flavobacteriaceae and *Prochlorococcus* across BBMO and MiCRO in relation to ENSO. **Panel A.** Abundance profile in Reads Per Kilobase per Million mapped reads (RPKM) through time of Flavobacteriaceae. **Panel B.** Abundance profile through time in Reads Per Kilobase per Million mapped reads (RPKM) of *Prochlorococcus*. Genome completeness (orange circle for BBMO and blue circle for MiCRO genomes) and ANI values (yellow circle) are shown in each panel. The Oceanic Niño Index (ONI), based on NOAA Niño 3.4 SST anomalies, is displayed as a blue dashed line on the secondary y-axis (right). Note that in MiCRO, Flavobacteriaceae abundance decreases during very strong El Niño events (e.g., 2015-2016), whereas *Prochlorococcus* shows the opposite trend, with marked increases during or just after these periods.

To further characterize microdiversity within these shared genomes, we assessed strain-level diversity by reconstructing haplotypes and clustering them into distinct strains for each time series. From the selected genome pairs, those belonging to SAR86 and Flavobacteriaceae yielded two strains per site in both BBMO and MiCRO (**Figure 5**). Strain dynamics remained relatively stable over time in both cases, with no clear dominance shifts or replacement events between strains for each genome. The Flavobacteriaceae genome in MiCRO, however, exhibited slightly greater fluctuations in relative abundance between the two strains than in BBMO (**Figure 5 E, F**). The two SAR86 genomes exhibited similar patterns in BBMO and MiCRO, with both strains coexisting consistently over time (**Figure 5 A-D**). Although strain inference was performed independently within each site, the genome pairs analyzed share high nucleotide identity (>98% ANI). Therefore, the strain clusters recovered within each microbiome may represent closely related subpopulations, but our analysis does not establish direct one-to-one correspondence of strains across sites. In specific genomes, distinct strains were detected at only one of the two sites (**Supplementary Figure S1**). These include the Ca. *Micropelagos* genome, in which two strains were identified in MiCRO, and Thalassobaculales, which also had 2 strains, but only in BBMO. In both cases, strain dynamics also remained stable over time, with two strains coexisting without clear replacement or persistent dominance shifts. Given its ecological relevance, contrasting abundance patterns, and high genome quality (BBMO long-read MAG: 77.9% completeness, 0.27% contamination; MiCRO SAG: 81.3% completeness, 0% contamination), we next focus on a *Prochlorococcus* genome pair (96% ANI), recovered as a single-amplified genome (SAG) in MiCRO and as a long-read MAG in BBMO, for further analyses.

**Figure 5.**
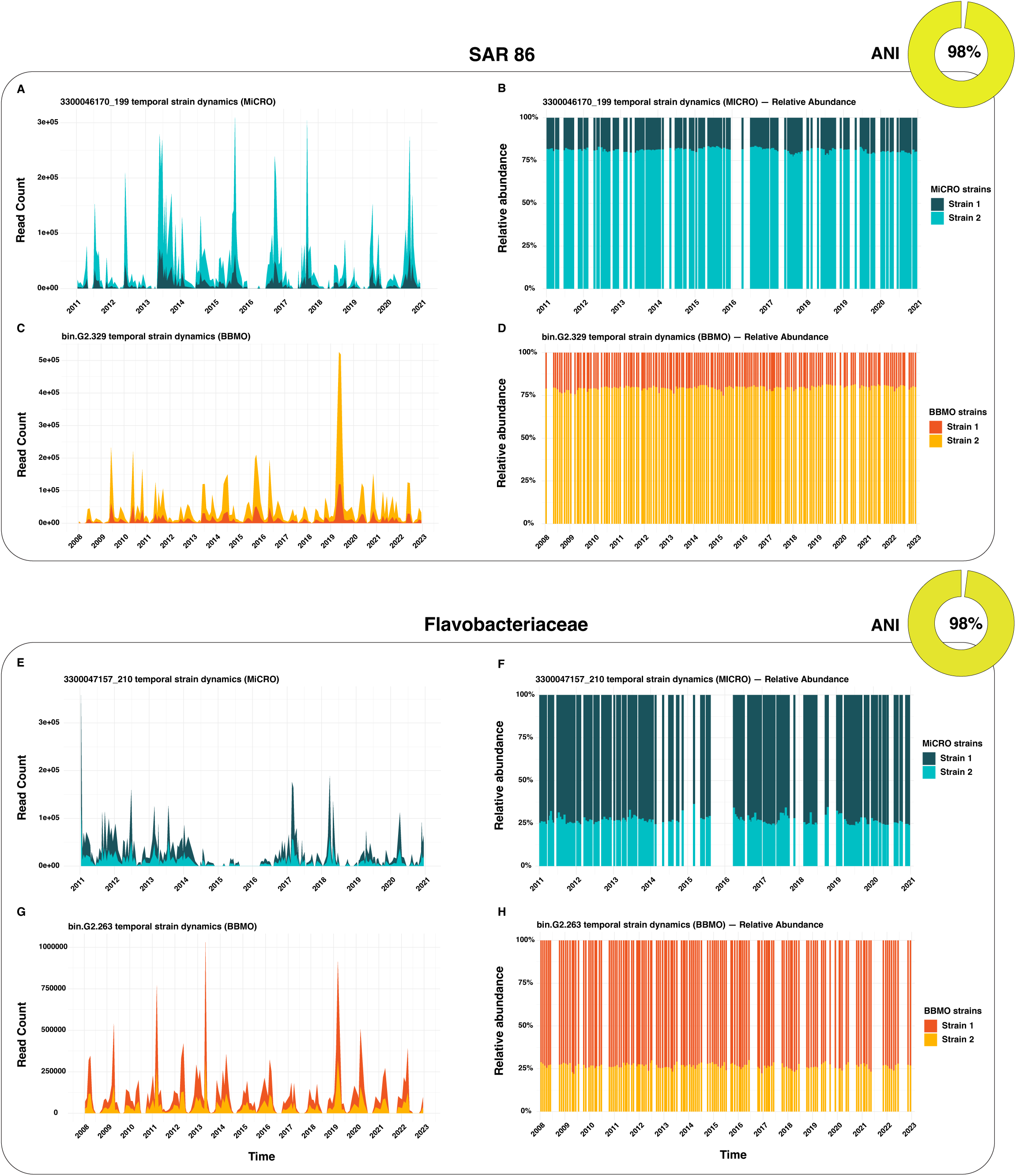
Strain-level dynamics of representative genomes present in BBMO and MiCRO. **Panels A and B.** Temporal strain dynamics in of MiCRO SAR86 genome in read counts and relative abundance respectively. **Panels C and D.** Temporal strain dynamics of BBMO SAR86 genome in read counts and relative abundance respectively. Both BBMO and MiCRO genome representatives share 98% ANI (yellow circle). **Panels E and F.** Temporal strain dynamics of MiCRO Flavobacteriaceae genome in read counts and relative abundance respectively. **Panels G and H.** Temporal strain dynamics of the BBMO Flavobacteriaceae genome in read counts and relative abundance, respectively. Both BBMO and MiCRO genome representatives share 98% ANI (yellow circle). BBMO genomes are shown in orange and MiCRO genomes in blue; Strain 1 is represented by the darker shade of each color, and Strain 2 by the lighter shade. Strain labels are defined independently within each site and do not imply correspondence between BBMO and MiCRO strains. These examples illustrate that shared genomes can harbor multiple strains that persist stably over decadal time scales, with limited evidence for strain replacement.

### *Prochlorococcus* genomes present in both locations

Whole-genome alignment revealed extensive synteny between the two *Prochlorococcus* genomes, with large co-linear regions and only a few structural rearrangements (**Figure 6A**), consistent with patterns previously observed in cultured *Prochlorococcus* genomes (*56*). Sequence identity across aligned regions ranged from 84% to 100%, with most homologous segments exceeding 94%. The 5S, 16S, and 23S ribosomal RNA genes were identified in both genomes with 97.11% sequence identity (**Figure 6A**), supporting a close phylogenetic relationship between the two *Prochlorococcus* genomes. Five local inversions were observed, involving conserved genes including *trpG* (tryptophan biosynthesis), *dgkA* (lipid metabolism), and *prfB* (translation termination) (**Figure 6A**, red ribbons). The BBMO long read partial genome consisted of 1,454 genes spanning 1.23 Mb across two contigs and is estimated to be 77.9% complete, corresponding to an estimated genome size of ∼1.57 Mb. In comparison, the MiCRO SAG contained 1,634 predicted genes across 1.43 Mb distributed over nine contigs with a completeness of 81.3%, yielding an estimated genome size of ∼1.76 Mb. Of these, 984 genes were present in the two genomes based on protein sequence homology, corresponding to 70.49% of the genes in the BBMO genome and 60.41% of the genes in the MiCRO genome (48% of the combined gene repertoire) (**Figure 6B**). The BBMO partial genome included 412 unique genes not found in the MiCRO genome, while the MiCRO genome contained 645 unique genes (**Figure 6B**). These differences can emerge from differences in genome content and completeness.

**Figure 6.**
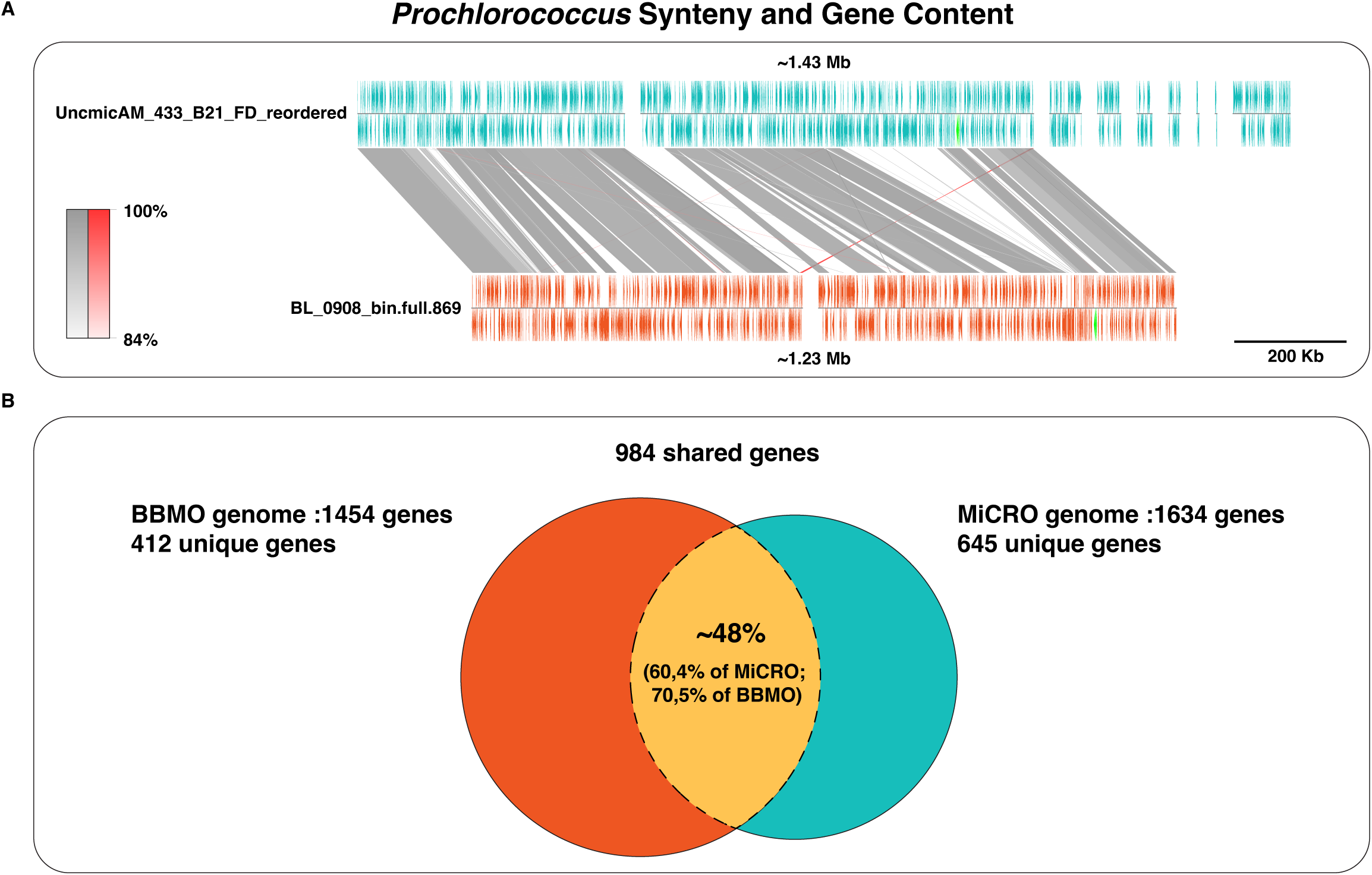
Genomic synteny and gene content comparison between *Prochlorococcus* present in both BBMO and MiCRO. **Panel A.** Partial-genome alignment of the BBMO long-read MAG in orange and the MiCRO SAG in blue (96% ANI) reveals extensive synteny, with large co-linear regions and five local inversions (red ribbons). Conserved ribosomal RNA operons (5S, 16S and 23S) are shown in green, with 97.11% sequence identity between genomes. Sequence identity between homologous regions is indicated by a grey scale (bottom left), and a 200 kb scale bar is shown for reference. **Panel B.** The partial BBMO genome in orange encodes 1,454 genes, and the MiCRO genome in blue 1,634 genes, of which 984 are shared (48% of the combined gene set). Genes uniquely detected in BBMO (n = 412) or MiCRO (n = 645) are indicated.

We further characterized the shared and unique genomic content of these genomes (**Figure S2**). Gene families were partitioned into persistent (generally considered core) and shell (variable) categories according to the PPanGGOLiN Pangenome framework, which defines the pangenome as the complete set of gene families observed across multiple *Prochlorococcus* genomes. All genes present in both *Prochlorococcus* genomes fell within the persistent partition, indicating a highly conserved core genome. A smaller fraction of gene families was assigned to the shell partition, representing genes present in only one genome. Additionally, the MiCRO genome displayed a higher proportion of single-copy genes, whereas the BBMO genome contained substantially more multicopy gene families (5 in MiCRO versus 46 in BBMO), contributing to differences in overall gene repertoire size between the two genomes.

### Genome-wide seasonal population structure and long-term dynamics

We assessed seasonal structure in population-level variant composition across genomes in both time series. For each genome, filtered genotype calls from the VCF files were converted into sample-by-variant presence/absence matrices, and seasonal differences were tested using PERMANOVA on Bray– Curtis dissimilarities. A significant seasonal effect was detected for most genomes in both MiCRO and BBMO, including 660 of 917 tested genomes in MiCRO (71.97%) and 957 of 1,451 in BBMO (65.95%). Similar proportions were observed in the shared dereplicated genome sets, where 98 of 124 genomes in MiCRO (79.03%) and 95 of 119 in BBMO (79.83%) also showed significant seasonal structure (**Table 2**). These results indicate that seasonal organization of within-population variant composition is a broad feature of microbial genomes in both ecosystems.

**Table 2.** Number of genomes showing significant seasonal structure of variants in MiCRO and BBMO across the full genome collection and the shared dereplicated genome set (95% ANI 70% Cov.).

| | *Seasonal effect*<br>(*PERMANOVA*, $p < 0.05$) | *No seasonal effect*<br>(*PERMANOVA*, $p \geq 0.05$) | *Excluded* | *Genomes tested* | *Total genomes* |
| --- | --- | --- | --- | --- | --- |
| *MiCRO* | 660 (71.97%) | 257 (28.03%) | 338 | 917 | 1255 |
| *MiCRO (dereplicated)* | 98 (79.03%) | 26 (20.97%) | 6 | 124 | 130 |
| *BBMO* | 957 (65.95%) | 494 (34.05%) | 84 | 1451 | 1535 |
| *BBMO (dereplicated)* | 95 (79.83%) | 24 (20.17%) | 1 | 119 | 120 |

We next tested whether interannual climatic variability associated with ENSO explained additional variation in population-level variant composition beyond seasonal structure. To this end, we used PERMANOVA models including both season and the Oceanic Niño Index (ONI), testing the marginal effect of ONI after accounting for season. As a result, we found a strong ENSO-associated structure in variant composition in MiCRO, in contrast to the much weaker signal detected in BBMO. In MiCRO, 677 of 890 tested genomes (76.06%) showed a significant ONI effect, whereas only 149 of 1,362 BBMO genomes (10.94%) did so. The same contrast was observed in the shared dereplicated genome set, where 94 of 123 MiCRO genomes (75.42%) showed significant ONI-associated structure compared with only 12 of 115 BBMO genomes (10.43%) (**Table 3**). These results indicate that, beyond seasonal organization, interannual ENSO-related variability is a major structuring force in MiCRO populations.

**Table 3.** ENSO-associated variation in genomic variant composition beyond seasonal variation across the full genome collection and the shared dereplicated genome set (95% ANI 70% Cov.).

| | *ENSO effect*<br>(*PERMANOVA*, $p < 0.05$) | *No ENSO effect*<br>(*PERMANOVA*, $p \geq 0.05$) | *Excluded* | *Genomes*<br>*tested* | *Total*<br>*genomes* |
| --- | --- | --- | --- | --- | --- |
| *MiCRO* | 677 (76.06%) | 213 (23.94%) | 365 | 890 | 1255 |
| *MiCRO (dereplicated)* | 94 (75.42%) | 29 (23.58%) | 7 | 123 | 130 |
| *BBMO* | 149 (10.94%) | 1213 (89.06%) | 173 | 1362 | 1535 |
| *BBMO (dereplicated)* | 12 (10.43%) | 103 (89.57%) | 5 | 115 | 120 |

We further explored these dynamics in the genetic composition of *Prochlorococcus* populations at MiCRO (SAG) and BBMO (long-read MAG) using a variant-based Principal Component Analysis (PCA). Also, in parallel, we assessed evolutionary signals by decomposing genome-wide pN/pS across time series **(Figure 7)**. Principal Component Analysis (PCA) of variant presence/absence matrices revealed that *Prochlorococcus* mutations exhibited seasonal signatures at both sites (**Figure 7A, B**). In these plots, each point represents a genetic variant or mutation detected in the Variant Calling Files (VCFs), colored by the season in which it was most frequently observed. Seasonal structuring of variants, particularly along the first principal component (PC1), indicates that different sets of mutations tend to dominate in different periods of the year, suggesting seasonally structured variation in the genetic composition of the *Prochlorococcus* populations. This pattern was consistent with the PERMANOVA results for both sites, which showed a significant effect of season on variant composition in BBMO (R^2^ = 0.152, adjusted p = 0.001) and MiCRO (R^2^ = 0.037, adjusted p = 0.001). Notably, the BBMO genome showed more dense seasonal aggregates of variants, especially in winter and spring, whereas the MiCRO *Prochlorococcus* genome appeared to have variants that were clustered into summer or autumn. A similar seasonal genetic signature was detected in other taxa beyond *Prochlorococcus*. For instance, a Flavobacteriaceae genome present in both locations exhibited significant seasonal structuring of variants in both BBMO (R^2^ = 0.180, adjusted p = 0.001) and MiCRO (R^2^ = 0.044, adjusted p = 0.001) (**Figure S3A, B**), reinforcing the idea that seasonality affects the structure of microbial populations.

**Figure 7.**
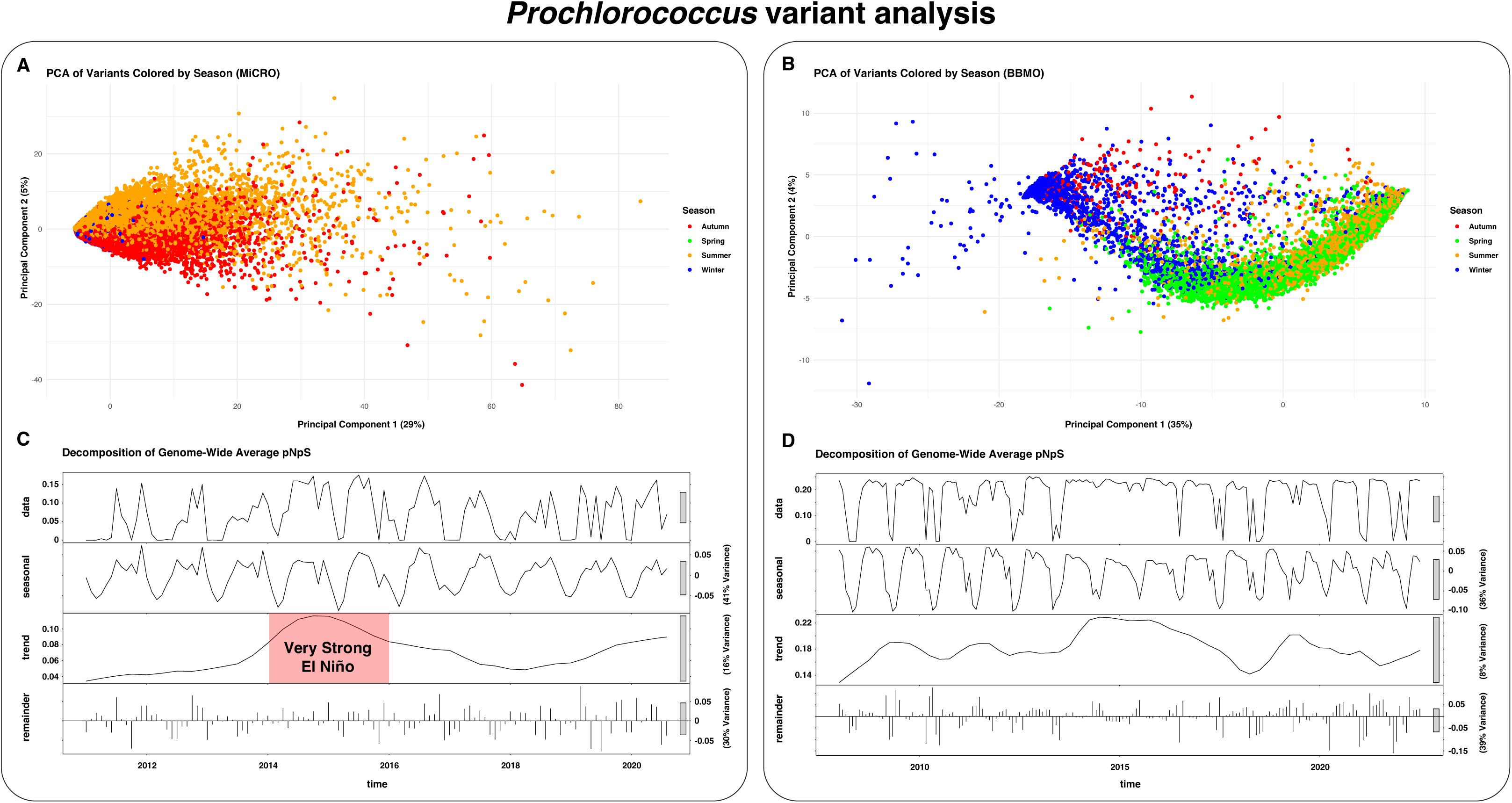
Seasonal and long-term population dynamics of shared *Prochlorococcus* genomes at BBMO and MiCRO. **Panels A and B.** Principal Component Analysis (PCA) of variants detected in the *Prochlorococcus* genomes from MiCRO (A) and BBMO (B). Each point represents a genetic variant or mutation detected in the corresponding VCFs after the variant calling analysis, colored according to the season where they appear the most (spring = green, summer = orange, autumn = red, winter = blue). Clustering of variants by season, especially along PC1 highlights cyclic seasonal shifts in the genetic composition of *Prochlorococcus* populations. Percent variance explained by each axis is indicated in parentheses. **Panels C and D.** Seasonal-Trend decomposition (STL) of the genome-wide average pN/pS ratio over time for the same *Prochlorococcus* genomes in MiCRO (C) and BBMO (D). For each site, the top panel (“data”) shows the observed genome-wide average pN/pS ratio in each time series, followed by the seasonal, trend, and remainder components. Seasonal panels capture cyclic annual fluctuations in selection; trend panels highlight long-term changes, with elevated values in MiCRO coinciding with strong El Niño periods (area labeled in red); and the remainder represents unexplained variance. The proportions of variance explained by the seasonal and trend components are indicated next to each panel. These analyses reveal that *Prochlorococcus* populations at both sites exhibit recurrent replacements, while MiCRO also shows changes in pN/pS associated with selection exerted by ENSO events.

To assess the adaptive signal over time, we analyzed the genome-wide average pN/pS ratio across the time series, which reflects the balance between non-synonymous and synonymous variation and provides a proxy for selective pressures acting on populations. We decomposed the pN/pS signal into seasonal, trend, and residual components using STL (Seasonal-Trend decomposition via LOESS) to separate recurrent seasonal patterns from longer-term signals. Both *Prochlorococcus* genomes from MiCRO and BBMO showed a clear seasonal component in pN/pS, reflecting annual fluctuations in the proportion of non-synonymous relative to synonymous mutations (**Figure 7C, D**). These patterns likely reflect recurring environmental selection pressures and population replacement in *Prochlorococcus*. In the MiCRO time series, seasonal fluctuations explained 41% of the variance in pN/pS, compared with 36% in BBMO. Seasonal components were also detected in the Flavobacteriaceae genome at both sites (13% variance explained in MiCRO and 59% in BBMO; **Figure S3C, D**).

In the *Prochlorococcus* genome from MiCRO, the trend component revealed a potential long-term signal that appeared to correspond to El Niño Southern Oscillation (ENSO) events (**Figure 7C, red ribbons, Figure 1C**) with 16% of pN/pS variance explained. During El Niño periods, highlighted by the red-shaded interval in the trend component between 2014 and 2016, pN/pS values increased, suggesting the continuous environmental selection of a differentially adapted variant in warmer ENSO waters. No evident signal was detected in the pN/pS trend component of BBMO, despite accounting for 8% of the variance (**Figure 7D**). The remainder (unexplained) variation accounted for 30% and 39% of the variance in MiCRO and BBMO, respectively. Interestingly, the MiCRO Flavobacteriaceae trend showed the opposite ENSO pattern compared to MiCRO *Prochlorococcus*, with a decrease in the trend component of the pN/pS signal during El Niño years (**Figure S3C**), highlighted by the red-shaded interval, explaining 32% of pN/pS variance. This is consistent with the concurrent decline in Flavobacteriaceae abundance during ENSO events (**Figure 4A**), which points to negative environmental selection. In BBMO Flavobacteriaceae, the trend component accounted for only 3% of the variance (**Figure S3D)**.

## DISCUSSION

Microbial populations are expected to respond rapidly to environmental heterogeneity, given their fine-grained adaptations to specific niches (*57*, *58*). Thus, they represent a sensitive lens for detecting shifts in ecosystem state. However, our understanding of seasonal changes in marine microbial populations in temperate regions, and whether these dynamics are similar or divergent across locations, remains limited. Similarly, the influence of large-scale climate oscillations on microbial populations remains poorly understood. Our decadal, strain-resolved population genomics analysis across two geographically distant coastal ecosystems (BBMO and MiCRO time series), provides a unique opportunity to examine: 1) how many populations from the same species are present at each location, 2) whether the same or similar microbial populations exhibit comparable seasonal cycles, and 3) whether longer-term climate variability, such as ENSO, also influences these dynamics. By reconstructing thousands of MAGs from short- and long-read sequencing, as well as SAGs, and tracking their abundance and genetic variation across 15 (BBMO) and 10 (MiCRO) years, we resolved population and strain-level dynamics at a fine-grained temporal scale.

Despite being located in distant and coastal basins, the semi-enclosed NW Mediterranean Sea (BBMO) and the open Pacific Ocean (MiCRO), both ecosystems are structured by strong seasonal cycles, including marked annual fluctuations in temperature (∼12-25 °C at BBMO and ∼12-22 °C at MiCRO) (*24*, *38*). However, in addition to this shared seasonal forcing, MiCRO is subject to marked interannual climatic variability driven by ENSO anomalies (*50*, *51*). To assess how microbial populations are structured under these environmental regimes, we clustered genomes retrieved from both locations using approximate species (95%) and strain (99%) ANI thresholds, widely used in microbial genomics (*41*, *42*), providing a framework for comparing population overlap across sites. Our genome reconstructions revealed a moderate overlap at the species-level: 250 genomes shared at 95% ANI out of 2,790 analyzed but a pronounced differentiation at the strain-level, with only 13 BBMO genomes dereplicating with 15 MiCRO genomes at 99% ANI. Thus, while both sites share several species and experience similar environmental heterogeneity, strain-level comparisons revealed that each microbiome harbors unique microdiversity, uncovering fine-grained biogeographic differences likely shaped by local conditions (*59*). Similar patterns of species-level ubiquity coupled with strain-level differentiation have been reported for specific marine microbial taxa. For example, SAR11 can be globally ubiquitous at the species level, yet exhibits geographically structured fine-scale genomic variation within subclades (*47*). Similarly, marine *Synechococcus* is organized into ecotypes with distinct environmental distributions rather than uniform global populations. For example, there are clades associated with colder, mesotrophic waters, others dominate warm oligotrophic open-ocean regions, and others are restricted to low-iron environments, indicating strong biogeographic partitioning across the global ocean (*60*). This underscores the importance of accounting for population-level variation to understand the spatiotemporal distributions and adaptations of marine microbes, and the environmental drivers shaping them.

The few strains that were present in both BBMO and MiCRO were preferentially detected in coastal Mediterranean climates. In contrast, shared genomes at the species-level were detected across multiple oceanic basins globally and spanned both coastal and open-ocean regions. Thus, increasing genome similarity revealed regionally constrained strains, with potential adaptations to coastal Mediterranean conditions and broad distributions within this climate zone. Comparable patterns have been reported in other microbial systems such as in hot springs, where closely related lineages recur across geographically distant yet environmentally similar habitats (*61*). This indicates that at least some ecotypes have a high dispersal capability and are able to reach their specific niches even if they are located far away. In addition, our results suggest that habitat filtering acts not only on taxonomic composition, but also on the genomic traits that favor persistence in particular environments. Cosmopolitan marine taxa adapted to oligotrophic open-ocean environments often exhibit streamlined genomes, characterized by reduced gene content and metabolic specialization, a pattern well documented for lineages such as SAR11 and *Prochlorococcus* (*54*, *62*). In contrast, microbes inhabiting coastal and temperate environments typically experience greater environmental variability in nutrient availability, temperature, and mixing regimes, and frequently harbor larger genomes with expanded functional repertoires (*55*), being able to grow fast when conditions are favorable (*55*). In this context, the genome size patterns we have observed align with the biogeographic structuring revealed by the genome similarity analysis. Genomes shared at the species level exhibited a broad global distribution and tended to be smaller, consistent with streamlined, cosmopolitan strategies. Their presence at both observatories may indicate that these locations are within their distribution range, or that they reached them via mass effects. In turn, the subset of strain-level genomes shared between BBMO and MiCRO exhibited slightly larger genome sizes and geographically restricted distributions, being preferentially detected in coastal environments, pointing to specific regional adaptations.

Despite the revealed strain-level differences between sites, several selected genomes showed strikingly seasonal and aligned across basins dynamics indicating shared adaptation. Previous long-term studies at individual coastal observatories have shown that marine microbial populations often exhibit recurrent seasonal abundance patterns driven by predictable environmental cycles (e.g., temperature, mixing, and nutrient availability) (*24*, *63*). Here, by directly comparing two distant coastal time series at population resolution, we show that such conserved phenologies can extend across ocean basins. High-quality representative genomes sharing *≥*96 ANI, such as Thalassobaculales and Ca. *Micropelagos*, showed recurrent summer/autumn maxima at both sites, consistent with conserved phenologies under a common annual cycle. In turn, whereas *Synechococcus* displayed sharp, high-amplitude peaks at BBMO and a more continuous and moderate presence in MiCRO, SAR86 exhibited a warm-season phase shift between sites. Together, these patterns suggest that broad seasonal forcing may organize abundance dynamics across distant coasts, while local factors modulate timing and magnitude (e.g., stratification, nutrient pulses, grazing/viral control) (*64*, *65*). This seasonal baseline provides the reference against which we assess departures linked to interannual climate variability at MiCRO. Building on this, interannual or long-term forcing at MiCRO revealed lineage-specific departures from these patterns. During strong El Niño events, Flavobacteriaceae abundance declined while *Prochlorococcus* abundance increased, suggesting that large-scale climatic anomalies can differentially affect microbial lineages within the same ecosystem. Similar shifts have been reported previously at the community and ecotype levels during El Niño, including enrichment of oligotrophic cyanobacteria and transitions from cold-to warm-adapted lineages (*38–40*). Our results indicate that such climate-linked responses are also evident at the population-genomic scale.

Time-series studies have reported genome-wide or gene-specific selective sweeps and lineage turnover under certain ecological contexts, particularly in freshwater bacterial populations and in marine archaeal nitrifiers experiencing strong or directional selection (*66*, *67*). Here, our strain-resolved analyses revealed that multiple coexisting strains persisted stably across the time series, including in MiCRO, with little evidence of replacement events in either microbiome. Despite pronounced fluctuations in overall abundance, these results indicate that population abundances can be highly sensitive to seasonal and interannual climatic variability without necessarily leading to strain replacement over decadal scales (*68*, *69*).

Across the broader genome set, most genomes (∼70%) showed significant seasonal structuring of variant composition. This indicates that seasonal organization of population genetic composition is widespread across both microbiomes, suggesting that recurrent annual environmental cycles structure microdiversity in a predictable manner. Beyond seasonality, our genome-wide analysis also showed that ENSO-associated variability explained additional structure in variant composition, particularly in MiCRO. This signal was widespread across MiCRO genomes (∼76%) but much weaker in BBMO (∼11%), consistent with the exposure of the MiCRO ecosystem to Pacific climate variability. To explore these patterns in greater detail across sites, we focused on a high-quality 96% ANI *Prochlorococcus* pair with representatives from both sites (a long-read MAG from BBMO and a SAG from MiCRO). Despite spanning distant ocean basins, the two genomes showed high synteny and extensive core gene conservation, with differences confined mainly to inversions and accessory gene content. Such patterns are consistent with prior work showing that *Prochlorococcus* genomes maintain a highly conserved backbone while varying in their flexible gene repertoires, which underlie ecological differentiation (*46*, *72*, *73*). The presence of large conserved blocks intercalated with site-specific accessory genes echoes observations from pangenomic analyses, in which gene gain, loss, and rearrangement sustain local adaptation within a globally cohesive lineage (*74–76*). The conserved genomic structure of the two genomes provided a solid basis for comparing population-genomic signals across oceans while recognizing bounded structural and gene-content differences that may tune lineage-specific responses. With this foundation, we next asked how selection shapes *Prochlorococcus* populations through time. Variant-based analyses revealed strong seasonal signatures at both sites, consistent with recurrent genetic turnover driven by annual environmental cycles (i.e., some variants are more frequent in winter, others in summer consistently). Similar seasonal variant clustering was observed in Flavobacteriaceae, indicating that this signature of seasonal structuring of variant composition extends across phylogenetically and ecologically distinct marine lineages. Consistently, Seasonal-Trend decomposition (STL) of genome-wide pN/pS ratios through time also revealed a strong seasonal component at both sites for these two genomes. The pN/pS ratio, which compares the frequency of nonsynonymous to synonymous mutations, provides a proxy for the balance between purifying (pN/pS <1) and adaptive (pN/pS >1) selection (*52*), underscoring that seasonal cycles modulate the genetic composition of microbial populations through time. Several freshwater studies have likewise documented seasonal shifts in strain composition over shorter time scales (*70*, *71*). A 20-year freshwater metagenomic time series showed that ∼80% of the microbial species showed cyclical, seasonal shifts in strain composition, supporting seasonality as a pervasive, cross-biome driver of population-genomic structure (*26*). That same study also reported that a smaller fraction of species (∼20%) underwent longer-term, decadal changes, pointing to additional layers of variability beyond the seasonal cycle. Extending beyond freshwater systems, here we detect such a long-term component at MiCRO, where the pN/pS trend in *Prochlorococcus* closely tracked ENSO anomalies, with elevated values during strong El Niño events, pointing to the persistent positive selection of a differentially adapted population during this event. Interestingly, this climate imprint was lineage-contingent, as Flavobacteriaceae exhibited the opposite association with ENSO, underscoring that large-scale climate variability can exert positive or negative selection across populations of microbial lineages. Together, these results point to a unifying mechanism across oceans: seasonal environmental selection establishes repeatable phenologies, while long-term climate modes (e.g., ENSO) represent an additional level at which selection acts, shaping the genomic composition of microbial populations. As recently suggested, ENSO-driven warming and associated nutrient limitation in coastal environments can produce ocean conditions similar to those predicted under climate change (*38*). In this context, recurrent ENSO anomalies may serve as natural experiments for understanding how marine microbial populations respond to future ocean warming and enhanced stratification.

## CONCLUSION

The comparative analysis of two long-term marine microbial observatories in contrasting coastal basins, the semi-enclosed NW Mediterranean Sea (BBMO) and the open Pacific Ocean (MiCRO), revealed that, despite similar seasonal environmental forcing and hundreds of shared species, strain-level assemblages remain largely distinct across basins, underscoring the importance of microdiversity in structuring marine populations. As genome similarity increased, shared genomes shifted from cosmopolitan toward regionally adapted populations characteristic of Mediterranean coastal environments. Across both ecosystems, population dynamics were strongly seasonal, with recurrent and cyclical shifts in variant composition consistent with annual changes in environmental selection. At the same time, ENSO anomalies at MiCRO introduced an additional level of selection, generating lineage-specific responses in opposite directions in *Prochlorococcus* and Flavobacteriaceae. These findings indicate that microbial populations, in specific locations, can be governed by a dual regime of seasonal recurrence and climate-associated selective shifts. More broadly, they suggest that understanding the dynamics of microbial populations may require accounting for both the seasonal selection cycles and the large-scale climate oscillations, which may also help anticipate microbial responses to future ocean change.

## MATERIALS AND METHODS

### Sampling and DNA extraction

Surface seawater samples (0.5 m depth) were collected monthly at the LTER Blanes Bay Microbial Observatory (BBMO; 41°40′N, 2°48′E; http://bbmo.icm.csic.es), located in the Bay of Blanes, Catalonia, Spain, between 2008 and 2023 (15 years). BBMO is an oligotrophic coastal site situated approximately 1 km offshore, with a water column depth of ∼20 meters and minimal influence from riverine or anthropogenic inputs. At each sampling event, ∼6 liters of surface seawater pre-filtered through a 200 μm mesh were sequentially filtered using a peristaltic pump through a 20 μm mesh, followed by a 3 μm pore-size polycarbonate filter (Poretics), and finally through a 0.22 μm polyethersulfone Sterivex filter (Millipore, Darmstadt, Germany) (*14*). For the MiCRO time series, surface seawater samples (0.5 m depth) were collected at Newport Pier (∼100 m from shoreline), Newport Beach, California, USA (33.608°N, 117.928°W), from 2011 to 2021 (10 years), at frequencies ranging from daily to monthly (averaged weekly). Four autoclaved bottles were rinsed with nearshore surface water before collection and transported immediately to the laboratory. For each sample, 1 liter of seawater was filtered sequentially through a 2.7 μm GF/D filter and a 0.22 μm Sterivex filter (Millipore, Darmstadt, Germany) using sterile tubing connected to a peristaltic pump. The filters were preserved with 1,620 μl of lysis buffer (23.4 mg/ml NaCl, 257 mg/ml sucrose, 50 mM Tris-HCl, 20 mM EDTA) and stored at −20 °C until extraction (*36*, *38*, *39*).

DNA extraction from BBMO samples followed the protocol described by Schauer et al. (*77*). Briefly, Sterivex cartridges were incubated with lysozyme solution (20 mg/ml) to lyse cells, followed by a second incubation with proteinase K (20 mg/ml). Lysates were pooled and extracted twice with phenol-chloroform-isoamyl alcohol (25:24:1, pH 8) and once with chloroform-isoamyl alcohol (24:1). The DNA was then concentrated using Amicon units (Millipore). Before sequencing, DNA quality was assessed via agarose gel electrophoresis, Qubit fluorometry, and Nanodrop spectrophotometry. DNA from MiCRO samples was extracted following an adapted protocol (*78*). Sterivex filters were first incubated at 37 °C for 30 minutes with lysozyme (final concentration 50 mg/ml). Proteinase K (1 mg/ml) and 10% SDS were then added, followed by overnight incubation at 55 °C. DNA was precipitated using ice-cold 100% isopropanol and sodium acetate (245 mg/ml, pH 5.2), pelleted by centrifugation, and resuspended in TE buffer (10 mM Tris-HCl, 1 mM EDTA) at 37 °C for 30 minutes. DNA was further purified using the Genomic DNA Clean and Concentrator kit (Zymo Research, Irvine, CA, USA) and stored at −80 °C. Before sequencing, DNA concentration was measured using the Qubit dsDNA HS assay kit and a Qubit fluorometer (Thermo Fisher Scientific, Waltham, MA, USA). DNA integrity was visually inspected via agarose gel electrophoresis, and purity was assessed with a NanoDrop ND-1000 spectrophotometer (Thermo Fisher Scientific, Waltham, MA, USA). Frozen DNA (25-500 μl in 1× TE buffer per sample) was then shipped to the DOE Joint Genome Institute (JGI) for sequencing.

### Shotgun metagenomic sequencing (short- and long-read)

The extracted DNA from BBMO was used for Illumina (short-read) and PacBio (long-read; Sequel II, HiFi) shotgun sequencing. For the short-read approach, a total of 174 samples collected between 2008 and 2023 (15 years) were processed. Specifically, samples collected from January 2009 to December 2011 (3 years) were sequenced using an Illumina HiSeq 4000 platform (2 × 150 bp), while samples from January to December 2008 (1 year) and from January 2012 to December 2023 (11 years) were sequenced on an Illumina NovaSeq 6000 platform (2 × 150 bp) at the Centre Nacional d’Anàlisi Genòmica (CNAG), Barcelona, Spain. In total, approximately 5.2 Tbp of short-read metagenomic data were generated across all years, averaging 30 Gbp per sample. For long-read sequencing at BBMO, three samples (February 2009, August 2009, and January 2010) were sequenced using the PacBio Sequel II platform. DNA was sheared to 12-16 kbp fragments, multiplexed, and then pooled, purified, and size-selected for fragments larger than 3 kbp. Sequencing was performed on an 8M SMRT cell with the Sequel II Binding Kit 2.0 and Sequencing Chemistry v2.0 (PacBio). HiFi circular consensus sequences (CCS) were generated using the CCS pipeline in SMRT Link (v10.1.0.119588) with default parameters. The resulting CCS reads were demultiplexed and assigned to individual samples. Sequencing was carried out at the Norwegian Sequencing Centre (www.sequencing.uio.no).

Likewise, the extracted DNA from MiCRO was used for Illumina (short-read) shotgun sequencing and PacBio (long-read; Sequel II, HiFi) shotgun sequencing. In the case of the short-read sequencing, a total of 240 samples collected between 2011 and 2021 (10 years) underwent sequencing at JGI. After quality control, 120 samples required additional size selection: 10 ng of genomic DNA was sheared to approximately 300 bp using an LE220-Plus Focused-ultrasonicator (Covaris) and purified with a double SPRI method using Mag-Bind Total Pure NGS beads (Omega Bio-Tek). Libraries were prepared with the KAPA HyperPrep kit (Roche) using one-tube end-repair, A-tailing, and ligation with NEXTFLEX UDI barcodes (PerkinElmer), followed by enrichment with seven cycles of PCR. Libraries were quantified with the KAPA Library Quantification Kit (Roche) on a Roche LightCycler 480 instrument. Sequencing was performed on an Illumina NovaSeq 6000 system using NovaSeq XP v1.5 reagent kits and S4 flow cells, with a 2 × 151 bp paired-end protocol. In total, 3.21 Tbp of short-read metagenomic data were generated. For the SAGs generation, individual cells were isolated via high-throughput single-cell isolation by Fluorescence-activated cell sorting (FACS), followed by cell lysis and whole-genome amplification as described by the Rinke protocol (*79*). Amplified DNA was then used to construct Illumina standard shotgun libraries, which were sequenced at JGI using the Illumina NovaSeq6000 S4 platform. Raw Illumina sequences were then quality filtered using BBTools (*80*), normalized using BBNorm (*80*) and error corrected using Tadpole (*80*). For long-read sequencing at MiCRO, DNA aliquots from eight sampling dates were sequenced using the PacBio Sequel II platform. These samples corresponded to four pre-El Niño years (January 12, 2011; April 27, 2011; October 19, 2011; and July 7, 2012) and four post-El Niño years (April 10, 2019; July 17, 2019; October 2, 2019; and January 8, 2020). DNA was subjected to additional quality control and size selection (BluePippin, Sage Science) prior to sequencing. Sequencing was carried out on the Sequel II system using HiFi circular consensus sequencing (CCS). For the April 27, 2011, sample, two sequencing runs were performed (SRA accessions SRR32280797 and SRR32280798), which were later combined during analysis.

### Co-Assembly and reconstruction of genomes

All BBMO short-read sequences were quality-filtered and adapter-trimmed using CUTADAPT v1.16 (*81*). Given the high computational demands of the full 15-year dataset, the delineation of BBMO short-read MAGs focused on a subset comprising 84 monthly metagenomes collected between January 2009 and December 2015 (7 years). To enable co-assembly of samples with similar community compositions, pairwise metagenomic similarities were estimated using Simka v1.5.2 (*82*), based on k-mer profiles (k = 21), a minimum Shannon index of 1.5, a minimum read length of 70 bp, and Bray-Curtis dissimilarities. The resulting distance matrix was hierarchically clustered in R (*83*) using UPGMA, defining four groups corresponding to seasonal periods: G1 (winter, 37 samples), G2 (spring, 9 samples), G3 (early summer, 14 samples), and G4 (late summer, 22 samples). Two samples were excluded as they did not cluster consistently with any defined group. Samples within each cluster were co-assembled using MEGAHIT v1.2.8 (*84*). To estimate contig abundances across samples prior to binning, the 84 BBMO metagenomes (2009–2015) were mapped back to the four cluster-specific co-assemblies with BWA v0.7.17-r1188 (*85*) in default mode. Samtools (*86*) v1.8 was used to filter out unmapped reads, secondary alignments, and alignments with mapping quality scores below 10 for G2, G3, and G4 and Samtools v1.12 for G1. MetaBAT v2.12.1 (*87*) was applied to the resulting BAM files using default settings and a minimum contig length of 2.5 kb to generate initial MAG sets for each group. Contig depth profiles from MetaBAT, along with the BAM files, were also provided to two additional binners, Concoct v0.4.2 (*88*) and MaxBin2 v2.2.5 (*89*), to produce complementary sets of MAGs, both run in default mode with a 2.5 kb minimum contig length. Outputs from all three binners were integrated and refined using MetaWrap v1.3 (*90*) in default mode. Only refined MAGs with an estimated completeness ≥50% and contamination ≤10% were retained. This process yielded a total of 2,311 short-read MAGs, distributed as follows: G1 (909 MAGs), G2 (347), G3 (457), and G4 (598). Taxonomic classification was performed with GTDB-Tk v1.5 (*91*) using the classify_wf workflow. To remove redundancy, all MAGs were dereplicated at 99% Average Nucleotide Identity (ANI) using dRep v2.3.2 (*92*), independently of their initial sample group. Ultimately, this resulted in a final set of 1,505 high-quality, non-redundant short-read MAGs.

For the reconstruction of BBMO long-read MAGs, individual sample assemblies as well as a combined co-assembly were generated using hifiasm-meta (*93*). From the single-sample assemblies, we recovered 20 single-contig MAGs, each featuring >50% estimated completeness and contig lengths exceeding 500 kb; no binning was required for these MAGs. To further expand MAG recovery, contigs from both the individual and co-assembled datasets were subjected to binning with the HiFi MAG pipeline v1.5 (https://github.com/PacificBiosciences/pb-metagenomics-tools/blob/master/docs/Tutorial-HiFi-MAG-Pipeline.md), which employs MetaBat2 (*94*). This approach yielded an additional 37 high-quality PacBio MAGs (completeness >70%, contamination <10%, and <20 contigs). All 57 resulting MAGs (from both single-contig and binned sources) were evaluated for completeness, contamination, strain heterogeneity, and redundancy using CheckM2 (*95*) and compareM (https://github.com/dparks1134/CompareM). MAGs sharing >99% genome-wide Average Amino Acid Identity (AAI) were considered redundant, and the highest-quality representative was selected using a quality score defined as completeness – 5 × contamination (*96*). Preference was given to binned MAGs only when they showed substantially higher quality than single-contig MAGs or when single-contig MAGs had completeness <70%. In cases where the quality was similar, single-contig MAGs were prioritized. Among bins with comparable quality and strain heterogeneity, those from individual sample assemblies were favored over co-assembly-derived bins. Following this curation process, a final set of 30 non-redundant long-read MAGs (<90% AAI) with completeness above 50% and contamination below 10% was retained for downstream analyses. Average nucleotide identity (ANI) was computed across the 30 MAGs using FastANI (*41*), confirming that all pairs exhibited ANI values below 95%, a commonly accepted species delineation threshold beyond which homologous recombination is minimal (*41*, *42*). At this threshold, the MAGs resolved into 30 independent species-level lineages.

For the co-assembly and reconstruction of MiCRO short-read MAGs, Illumina metagenomes from MiCRO were processed with the NMDC standardized workflows (*97*). Raw reads underwent adapter/contaminant trimming and decontamination with the Reads QC Workflow (v1.0.14-alpha.1), which uses BBDuk (BBTools v39.03) (*98*) for quality trimming, adapter/linker removal, spike-in removal, and host screening following NMDC defaults. Next, the corrected reads were assembled using metaSPAdes v4.0.0 (*99*); assemblies were validated and read coverage estimated with BBMap (*98*) using default workflow parameters. For genome binning, we applied the Metagenome Assembled Genomes (MAGs) Workflow (v1.3.14), which computes contig depth profiles from read mappings and bins contigs using MetaBAT 2 v2.15. (*94*) MAG quality (completeness/contamination) was estimated with CheckM v1.2.1 (*100*) and reported in the context of MiMAG standards using annotated rRNAs/tRNAs and single-copy marker genes; taxonomic assignment followed GTDB-Tk v2.1.1 (*91*) defaults as implemented in the NMDC MAGs workflow. For the reconstruction of the MiCRO short-read SAGs, artifact-filtered and normalized Illumina reads were assembled with SPAdes (v3.15.3) (*101*). Subsequently, 200 bp were trimmed from both ends of all contigs, and contigs shorter than 2 kb or with read coverage below 2 were discarded. Also, assemblies with a total size smaller than 20 kb were excluded.

PacBio HiFi MiCRO metagenomes were assembled with the long-read branch of the NMDC Metagenome Assembly Workflow (v1.0.7) (*97*), which uses Flye v2.9.2 (*102*) for assembly, pbmm2 (https://github.com/PacificBiosciences/pbmm2) for long-read alignment, Racon v1.4.20 (*103*) for polishing, and minimap2 v2:2.25 (*104*) for read mapping and coverage estimation using default parameters. Resulting long-read assemblies were binned with MetaBAT 2 v2.15 (*94*) via the MAGs Workflow (v1.3.14); bin quality and MiMAG reporting followed the same procedure as for short-read MAGs (CheckM v1.2.1, rRNAs/tRNAs/marker genes, GTDB-Tk v2.1.1). To remove redundancy and ensure consistency with the BBMO genome set, MiCRO MAGs and SAGs were dereplicated at 99% Average Nucleotide Identity (ANI) using dRep v2.3.2 (*92*), resulting in a final MiCRO genome set comprising 1,068 MAGs and 187 SAGs.

### Abundance and Horizontal coverage across samples

For downstream analyses, we focused on a total of 1,535 high-quality MAGs from BBMO (including both short- and long-read assemblies), 1,068 MAGs from MiCRO (also including short- and long-read assemblies), and 187 SAGs from MiCRO. To estimate genome abundances consistently across all datasets and minimize the number of incorrectly mapped reads, we used a competitive mapping strategy. Specifically, all 2,790 genomes were concatenated into two composite FASTA files, one for BBMO (1,535 genomes) and one for MiCRO (1,255 genomes). Clean reads from each site were then mapped against their respective concatenated genome file using BWA-MEM2 v2.2.1 (*85*), retaining only alignments with identity >95% and read alignment coverage >80%. From the resulting mappings, individual BAM files for each genome and sample were extracted with Samtools v1.8 (*105*). Finally, RPKM values (reads per kilobase of genome per million mapped reads) and horizontal genomic coverage (percentage of the genome covered by at least one filtered read) were computed for each genome in each metagenomic sample using CoverM v0.7.0 (*106*).

### Variant calling

To investigate genetic divergence of the BBMO and MiCRO genomes through time, we performed variant calling to identify nucleotide polymorphisms (SNPs), insertions, and deletions in all reconstructed genomes. For each genome and dataset, we first merged all individual BAM files into a single file using Samtools v1.8. The resulting merged BAM files were then processed with Freebayes v1.3.1 (*107*), setting ploidy to 1 (-p 1) and requiring at least four supporting observations for an alternate allele (-C 4). The resulting variant call format (VCF) files were subsequently analyzed with POGENOM v0.8.473 (*108*), using a minimum locus coverage of ten (--min_count 10) and requiring presence in at least four samples (--min_found 4). This analysis enabled us to compute the non-synonymous to synonymous mutation ratio (pN/pS) for each predicted gene across samples in all 2,790 genomes. We resolved haplotypes and quantified strain dynamics within each genome using Floria (v1.0) (*109*) and its companion tool Floria–Strainer (*110*). Floria reconstructs microbial haplotypes by solving a minimum error correction (MEC) problem, integrating information from BAM alignments, VCF variant calls, and the genome FASTA sequence. This process generates “haplosets,” which are groups of reads representing distinct strain-specific haplotypes. Floria–Strainer (in split mode) then clusters these haplosets into discrete strains using a Gaussian Mixture Model based on allele-frequency patterns, producing separate BAM files for each strain. Finally, each strain-specific BAM was processed with CoverM (v0.7.0) (*106*) to calculate RPKM and horizontal genome coverage, allowing us to quantify and track strain dynamics across the time series.

### Identification of genomes present in both time series

To identify genomes present in both microbiomes, a dereplication was performed using dRep (92). We delimited 95% ANI as the species-level threshold and 99% ANI as the strain-level threshold according to the literature (41). This analysis allowed us to identify genome clusters shared between the two time series at both taxonomic resolutions and to quantify the extent of genome overlap between sites. Among the genome pairs recovered through this dataset-wide comparison, we selected a Prochlorococcus pair sharing >96% ANI, which corresponds to a long-read MAG from BBMO (77.9% completeness, 0.27% contamination) and a SAG from MiCRO (81.3% completeness, 0% contamination), because of its ecological relevance and relatively high quality. We first annotated both BBMO and MiCRO genomes using Prokka (111) to generate General Feature Format (GFF) files, which were then used in downstream analyses. Using pyGenomeViz (112), we examined genomic synteny in both genomes. Additionally, we employed PPanGGOLiN (113) to analyze the pangenome, which categorizes genes as core (persistent, shared across all genomes) and shell (genes present in several but not all genomes).

To further characterize the ecological dynamics of dereplicated genomes, we examined abundance profiles (RPKM) through time for genomes shared between BBMO and MiCRO at minimum 95% ANI. To illustrate these general patterns, we show abundance time series for a representative subset of six dereplicated genomes (**Figure 3**), selected based on their elevated completeness, relatively high abundance across the time series, high ANI and ecological relevance, thereby providing representative snapshots of how genome abundances vary across distant coastal ecosystems.

### Statistics

Data analyses were performed using Python v3.8.5 (*114*) and R v3.5 (*83*). To investigate seasonal patterns in variant composition of genomes present in both locations, we analyzed VCF files by extracting genotype (GT) matrices using the R package vcfR (*115*), retaining only variants with a quality score >20. Genotypes were converted to a binary format, where 0 indicated reference and any non-zero value indicated the presence of a variant. Variants were further filtered to retain those present in more than 5% and less than 90% of samples. Samples were assigned to seasons according to their collection dates. For each genome, seasonal differences in sample-level variant composition were tested using PERMANOVA with 999 permutations on Bray-Curtis dissimilarities, as implemented in the R package vegan (*116*). To further visualize seasonal structure in representative genomes present at both locations, including *Prochlorococcus* and Flavobacteriaceae, we applied Principal Component Analysis (PCA) to binary variant matrices. Each variant was labelled according to the season in which it was most frequently detected, allowing seasonal clustering of variants to be visualized in PCA space.

To analyze the temporal structure of the pN/pS signal of the *Prochlorococcus* pair of genomes present in both sites in detail and decompose it into its fundamental components, we used Seasonal-Trend decomposition using LOESS (STL), implemented in R’s stats package (v3.6.2). STL is a statistical method that separates a time series into three principal components: seasonal component, capturing recurrent periodic patterns (e.g., annual cycles due to temperature or nutrient changes); trend component, reflecting long-term, gradual changes (e.g., driven by global climate forces or ecosystem shifts) and residual component, which includes unexplained variability, random noise, or anomalies. Applying STL to our pN/pS time series allowed us to separate the recurring seasonal variation from the long-term trend component. All data visualization was performed with *ggplot2* v3.2.

## ACKNOWLEDGMENTS

We would like to acknowledge all the members of the BBMO sampling team (http://bbmo.icm.csic.es), the MiCRO sampling team and the Ecology of Marine Microbes group (http://emm.icm.csic.es) and would like to thank the MARBITS platform of the Institut de Ciencies del Mar (ICM; http://marbits.icm.csic.es), the Finisterrae III supercomputer at the Centro de Supercomputación de Galicia (CESGA; https://www.cesga.es/) and the High Performance Community Computing Cluster (HPC3; https://rcic.uci.edu/hpc3/hpc3.html) for supporting the bioinformatics analyses. This paper acknowledges the generic ‘Severo Ochoa Centre of Excellence’ accreditation (CEX2019-000928-S).

## Funding

This investigation was supported by the project MINIME (PID2019-105775RB-I00, MICINN, Spain) and MAORI (PID2022-136281NB-I00, MICINN, Spain) to R.L.

## Author contributions

S.G.M. performed the statistical and bioinformatic analyses, including genome dereplication and variant analysis, contributed to their design and drafted the original manuscript. L.M. contributed to computational analyses related to BBMO time series. A.A.L. contributed to the generation and curation of MiCRO time series data. F.L conducted computational analyses including reconstruction and curation of the short-read BBMO MAGs. C.R.G carried out sample collection at BBMO and performed DNA extractions. T.H contributed to the reconstruction of BBMO long-read genomes. V.B. carried out sample collection at BBMO and performed DNA extractions. J.G. contributed to the generation of BBMO dataset. A.C.M provided MiCRO time series data and contributed to the analysis design. R.L. conceived the project, secured funding, designed the overall analytical strategy, and contributed to manuscript writing. All authors revised the manuscript and approved the final version.

## Competing interests

The authors declare no competing interests.

## Data and materials availability

LTER Blanes Bay Microbial Observatory (BBMO) short-read metagenomic sequences and associated contextual metadata are publicly available at the European Nucleotide Archive (ENA) under accession numbers <u>PRJEB48035</u> and <u>PRJEB51979</u>. PacBio Sequel II HiFi long-read sequences generated for BBMO are available at the ENA under accession number PRJEB106312. Short-read metagenome-assembled genomes (MAGs) reconstructed from the BBMO time series are publicly available at Zenodo (https://doi.org/10.5281/zenodo.17159964). This repository contains the first version (v1) of BBMO MAGs, including genome sequences, functional annotations, and abundance tables. *Tara* Oceans (TARA) abundances of MAGs are available at (https://doi.org/10.5281/zenodo.17634504). PacBio long-read MAGs reconstructed from BBMO are publicly available at Zenodo (https://doi.org/10.5281/zenodo.21517772).

Microbes in the Coastal Region of Orange County (MiCRO) short-read metagenomic sequences and associated contextual metadata are publicly available through the National Center for Biotechnology Information Sequence Read Archive (BioProject IDs: PRJNA1220992, PRJNA624320).

Custom scripts used for data processing and analysis are available from the corresponding author upon request.

## Notes

https://www.ebi.ac.uk/ena/browser/view/PRJEB48035

https://www.ebi.ac.uk/ena/browser/view/PRJEB51979

https://zenodo.org/records/17159964

https://zenodo.org/records/17634504

https://www.ncbi.nlm.nih.gov/bioproject/PRJNA1220992/

https://www.ncbi.nlm.nih.gov/bioproject/?term=PRJNA624320

https://doi.org/10.5281/zenodo.21517772

### Competing Interest Statement

The authors have declared no competing interest.

## REFERENCES

1. P. Falkowski, Ocean Science: The power of plankton. Nature 483, S17–20 (2012).

2. P. G. Falkowski, T. Fenchel, E. F. Delong, The Microbial Engines That Drive Earth’s Biogeochemical Cycles. Science 320, 1034–1039 (2008).

3. A. Z. Worden, M. J. Follows, S. J. Giovannoni, S. Wilken, A. E. Zimmerman, P. J. Keeling, Rethinking the marine carbon cycle: Factoring in the multifarious lifestyles of microbes. Science 347, 1257594 (2015).

4. P. A. del Giorgio, C. M. Duarte, Respiration in the open ocean. Nature 420, 379–384 (2002).

5. F. Azam, F. Malfatti, Microbial structuring of marine ecosystems. Nat Rev Microbiol 5, 782–791 (2007).

6. R. Logares, Decoding populations in the ocean microbiome. Microbiome 12, 67 (2024).

7. T. Van Rossum, P. Ferretti, O. M. Maistrenko, P. Bork, Diversity within species: interpreting strains in microbiomes. Nat Rev Microbiol 18, 491–506 (2020).

8. R. Logares, A. Boltovskoy, S. Bensch, J. Laybourn-Parry, K. Rengefors, Genetic diversity patterns in five protist species occurring in lakes. Protist 160, 301–317 (2009).

9. M. Schloter, M. Lebuhn, T. Heulin, A. Hartmann, Ecology and evolution of bacterial microdiversity. FEMS Microbiol Rev 24, 647–660 (2000).

10. G. L. Brennan, R. Logares, Tracking contemporary microbial evolution in a changing ocean. Trends Microbiol 31, 336–345 (2023).

11. S. Sunagawa, L. P. Coelho, S. Chaffron, J. R. Kultima, K. Labadie, G. Salazar, B. Djahanschiri, G. Zeller, D. R. Mende, A. Alberti, F. M. Cornejo-Castillo, P. I. Costea, C. Cruaud, F. d’Ovidio, S. Engelen, I. Ferrera, J. M. Gasol, L. Guidi, F. Hildebrand, F. Kokoszka, C. Lepoivre, G. Lima-Mendez, J. Poulain, B. T. Poulos, M. Royo-Llonch, H. Sarmento, S. Vieira-Silva, C. Dimier, M. Picheral, S. Searson, S. Kandels-Lewis, Tara Oceans coordinators, C. Bowler, C. de Vargas, G. Gorsky, N. Grimsley, P. Hingamp, D. Iudicone, O. Jaillon, F. Not, H. Ogata, S. Pesant, S. Speich, L. Stemmann, M. B. Sullivan, J. Weissenbach, P. Wincker, E. Karsenti, J. Raes, S. G. Acinas, P. Bork, E. Boss, C. Bowler, M. Follows, L. Karp-Boss, U. Krzic, E. G. Reynaud, C. Sardet, M. Sieracki, D. Velayoudon, Structure and function of the global ocean microbiome. Science 348, 1261359 (2015).

12. G. Salazar, L. Paoli, A. Alberti, J. Huerta-Cepas, H.-J. Ruscheweyh, M. Cuenca, C. M. Field, L. P. Coelho, C. Cruaud, S. Engelen, A. C. Gregory, K. Labadie, C. Marec, E. Pelletier, M. Royo-Llonch, S. Roux, P. Sánchez, H. Uehara, A. A. Zayed, G. Zeller, M. Carmichael, C. Dimier, J. Ferland, S. Kandels, M. Picheral, S. Pisarev, J. Poulain, S. G. Acinas, M. Babin, P. Bork, C. Bowler, C. de Vargas, L. Guidi, P. Hingamp, D. Iudicone, L. Karp-Boss, E. Karsenti, H. Ogata, S. Pesant, S. Speich, M. B. Sullivan, P. Wincker, S. Sunagawa, S. G. Acinas, M. Babin, P. Bork, E. Boss, C. Bowler, G. Cochrane, C. de Vargas, M. Follows, G. Gorsky, N. Grimsley, L. Guidi, P. Hingamp, D. Iudicone, O. Jaillon, S. Kandels-Lewis, L. Karp-Boss, E. Karsenti, F. Not, H. Ogata, S. Pesant, N. Poulton, J. Raes, C. Sardet, S. Speich, L. Stemmann, M. B. Sullivan, S. Sunagawa, P. Wincker, Gene Expression Changes and Community Turnover Differentially Shape the Global Ocean Metatranscriptome. Cell 179, 1068–1083.e21 (2019).

13. J. A. Cram, C.-E. T. Chow, R. Sachdeva, D. M. Needham, A. E. Parada, J. A. Steele, J. A. Fuhrman, Seasonal and interannual variability of the marine bacterioplankton community throughout the water column over ten years. ISME J 9, 563–580 (2015).

14. A. Auladell, I. Ferrera, L. Montiel Fontanet, C. D. Santos Júnior, M. Sebastián, R. Logares, J. M. Gasol, Seasonality of biogeochemically relevant microbial genes in a coastal ocean microbiome. Environ Microbiol, doi: 10.1111/1462-2920.16367 (2023).

15. S. Lambert, M. Tragin, J.-C. Lozano, J.-F. Ghiglione, D. Vaulot, F.-Y. Bouget, P. E. Galand, Rhythmicity of coastal marine picoeukaryotes, bacteria and archaea despite irregular environmental perturbations. ISME J 13, 388–401 (2019).

16. O. P. Rajora, Ed., Population Genomics: Concepts, Approaches and Applications (Springer International Publishing, Cham, 2019; https://link.springer.com/10.1007/978-3-030-04589-0)*Population Genomics*.

17. D. Garant, S. E. Forde, A. P. Hendry, The multifarious effects of dispersal and gene flow on contemporary adaptation. Functional Ecology 21, 434–443 (2007).

18. J. A. Fuhrman, J. A. Cram, D. M. Needham, Marine microbial community dynamics and their ecological interpretation. Nat Rev Microbiol 13, 133–146 (2015).

19. J. A. Fuhrman, I. Hewson, M. S. Schwalbach, J. A. Steele, M. V. Brown, S. Naeem, Annually reoccurring bacterial communities are predictable from ocean conditions. Proceedings of the National Academy of Sciences 103, 13104–13109 (2006).

20. J. A. Gilbert, D. Field, P. Swift, L. Newbold, A. Oliver, T. Smyth, P. J. Somerfield, S. Huse, I. Joint, The seasonal structure of microbial communities in the Western English Channel. Environmental Microbiology 11, 3132–3139 (2009).

21. E. J. Raes, S. Myles, L. MacNeil, M. Wietz, C. Bienhold, K. Tait, P. J. Somerfield, A. Bissett, J. van de Kamp, J. M. Gasol, R. Massana, Y.-C. Yeh, J. A. Fuhrman, J. LaRoche, Seasonal patterns of microbial diversity across the world oceans. Limnology and Oceanography Letters 9, 512– 523 (2024).

22. L. Alonso-Sáez, V. Balagué, E. L. Sà, O. Sánchez, J. M. González, J. Pinhassi, R. Massana, J. Pernthaler, C. Pedrós-Alió, J. M. Gasol, Seasonality in bacterial diversity in north-west Mediterranean coastal waters: assessment through clone libraries, fingerprinting and FISH. FEMS Microbiol Ecol 60, 98–112 (2007).

23. J. A. Gilbert, J. A. Steele, J. G. Caporaso, L. Steinbrück, J. Reeder, B. Temperton, S. Huse, A. C. McHardy, R. Knight, I. Joint, P. Somerfield, J. A. Fuhrman, D. Field, Defining seasonal marine microbial community dynamics. ISME J 6, 298–308 (2012).

24. A. Auladell, A. Barberán, R. Logares, E. Garcés, J. M. Gasol, I. Ferrera, Seasonal niche differentiation among closely related marine bacteria. ISME J 16, 178–189 (2022).

25. I. M. Deutschmann, A. K. Krabberød, F. Latorre, E. Delage, C. Marrasé, V. Balagué, J. M. Gasol, R. Massana, D. Eveillard, S. Chaffron, R. Logares, Disentangling temporal associations in marine microbial networks. Microbiome 11, 83 (2023).

26. R. R. Rohwer, M. Kirkpatrick, S. L. Garcia, M. Kellom, K. D. McMahon, B. J. Baker, Two decades of bacterial ecology and evolution in a freshwater lake. Nat Microbiol 10, 246–257 (2025).

27. K. L. Vergin, B. Beszteri, A. Monier, J. C. Thrash, B. Temperton, A. H. Treusch, F. Kilpert, A. Z. Worden, S. J. Giovannoni, High-resolution SAR11 ecotype dynamics at the Bermuda Atlantic Time-series Study site by phylogenetic placement of pyrosequences. ISME J 7, 1322–1332 (2013).

28. V. Tai, B. Palenik, Temporal variation of Synechococcus clades at a coastal Pacific Ocean monitoring site. ISME J 3, 903–915 (2009).

29. B. M. Robicheau, J. Tolman, D. Desai, J. LaRoche, Microevolutionary patterns in ecotypes of the symbiotic cyanobacterium UCYN-A revealed from a Northwest Atlantic coastal time series. Science Advances 9, eadh9768 (2023).

30. B. B. Tolar, L. Reji, J. M. Smith, M. Blum, J. T. Pennington, F. P. Chavez, C. A. Francis, Time series assessment of Thaumarchaeota ecotypes in Monterey Bay reveals the importance of water column position in predicting distribution–environment relationships. Limnology and Oceanography 65, 2041–2055 (2020).

31. J. C. Robidart, C. M. Preston, R. W. Paerl, K. A. Turk, A. C. Mosier, C. A. Francis, C. A. Scholin, J. P. Zehr, Seasonal Synechococcus and Thaumarchaeal population dynamics examined with high resolution with remote in situ instrumentation. ISME J 6, 513–523 (2012).

32. Z. Liu, M. Alexander, Atmospheric bridge, oceanic tunnel, and global climatic teleconnections. Reviews of Geophysics 45 (2007).

33. M. A. Cane, Oceanographic Events During El Niño. Science 222, 1189–1195 (1983).

34. M. J. McPhaden, X. Yu, Equatorial waves and the 1997–98 El Niño. Geophysical Research Letters 26, 2961–2964 (1999).

35. M. G. Jacox, E. L. Hazen, K. D. Zaba, D. L. Rudnick, C. A. Edwards, A. M. Moore, S. J. Bograd, Impacts of the 2015–2016 El Niño on the California Current System: Early assessment and comparison to past events. Geophysical Research Letters 43, 7072–7080 (2016).

36. A. C. Martiny, A. Talarmin, C. Mouginot, J. A. Lee, J. S. Huang, A. G. Gellene, D. A. Caron, Biogeochemical interactions control a temporal succession in the elemental composition of marine communities. Limnology and Oceanography 61, 531–542 (2016).

37. L. E. Lilly, M. D. Ohman, CCE IV: El Niño-related zooplankton variability in the southern California Current System. Deep Sea Research Part I: Oceanographic Research Papers 140, 36–51 (2018).

38. A. A. Larkin, M. L. Brock, A. J. Fagan, A. R. Moreno, S. D. Gerace, L. E. Lees, S. A. Suarez, E. A. Eloe-Fadrosh, A. Martiny, Climate-driven succession in marine microbiome biodiversity and biogeochemical function. Res Sq, rs.3.rs-4682733 (2024).

39. A. A. Larkin, A. R. Moreno, A. J. Fagan, A. Fowlds, A. Ruiz, A. C. Martiny, Persistent El Niño driven shifts in marine cyanobacteria populations. PLoS One 15, e0238405 (2020).

40. Y.-C. Yeh, J. A. Fuhrman, Effects of phytoplankton, viral communities, and warming on free-living and particle-associated marine prokaryotic community structure. Nat Commun 13, 7905 (2022).

41. C. Jain, L. M. Rodriguez-R, A. M. Phillippy, K. T. Konstantinidis, S. Aluru, High throughput ANI analysis of 90K prokaryotic genomes reveals clear species boundaries. Nat Commun 9, 5114 (2018).

42. M. R. Olm, A. Crits-Christoph, S. Diamond, A. Lavy, P. B. Matheus Carnevali, J. F. Banfield, Consistent Metagenome-Derived Metrics Verify and Delineate Bacterial Species Boundaries. mSystems 5, e00731–19 (2020).

43. A. M. Simpson, A. B. Chase, A. Rodríguez-Verdugo, J. B. Martiny, Investigating bacterial evolution in nature with metagenomics. Current Opinion in Microbiology 87, 102654 (2025).

44. P. Flombaum, J. L. Gallegos, R. A. Gordillo, J. Rincón, L. L. Zabala, N. Jiao, D. M. Karl, W. K. W. Li, M. W. Lomas, D. Veneziano, C. S. Vera, J. A. Vrugt, A. C. Martiny, Present and future global distributions of the marine Cyanobacteria Prochlorococcus and Synechococcus. Proceedings of the National Academy of Sciences 110, 9824–9829 (2013).

45. R. M. Morris, M. S. Rappé, S. A. Connon, K. L. Vergin, W. A. Siebold, C. A. Carlson, S. J. Giovannoni, SAR11 clade dominates ocean surface bacterioplankton communities. Nature 420, 806–810 (2002).

46. N. Kashtan, S. E. Roggensack, S. Rodrigue, J. W. Thompson, S. J. Biller, A. Coe, H. Ding, P. Marttinen, R. R. Malmstrom, R. Stocker, M. J. Follows, R. Stepanauskas, S. W. Chisholm, Single-cell genomics reveals hundreds of coexisting subpopulations in wild Prochlorococcus. Science 344, 416–420 (2014).

47. T. O. Delmont, E. Kiefl, O. Kilinc, O. C. Esen, I. Uysal, M. S. Rappé, S. Giovannoni, A. M. Eren, Single-amino acid variants reveal evolutionary processes that shape the biogeography of a global SAR11 subclade. eLife 8, e46497 (2019).

48. A. C. Gregory, A. A. Zayed, N. Conceição-Neto, B. Temperton, B. Bolduc, A. Alberti, M. Ardyna, K. Arkhipova, M. Carmichael, C. Cruaud, C. Dimier, G. Domínguez-Huerta, J. Ferland, S. Kandels, Y. Liu, C. Marec, S. Pesant, M. Picheral, S. Pisarev, J. Poulain, J.-É. Tremblay, D. Vik, S. G. Acinas, M. Babin, P. Bork, E. Boss, C. Bowler, G. Cochrane, C. de Vargas, M. Follows, G. Gorsky, N. Grimsley, L. Guidi, P. Hingamp, D. Iudicone, O. Jaillon, S. Kandels-Lewis, L. Karp-Boss, E. Karsenti, F. Not, H. Ogata, S. Pesant, N. Poulton, J. Raes, C. Sardet, S. Speich, L. Stemmann, M. B. Sullivan, S. Sunagawa, P. Wincker, M. Babin, C. Bowler, A. I. Culley, C. de Vargas, B. E. Dutilh, D. Iudicone, L. Karp-Boss, S. Roux, S. Sunagawa, P. Wincker, M. B. Sullivan, Marine DNA Viral Macro- and Microdiversity from Pole to Pole. Cell 177, 1109–1123.e14 (2019).

49. T. O. Delmont, M. Gaia, D. D. Hinsinger, P. Frémont, C. Vanni, A. Fernandez-Guerra, A. M. Eren, A. Kourlaiev, L. d’Agata, Q. Clayssen, E. Villar, K. Labadie, C. Cruaud, J. Poulain, C. Da Silva, M. Wessner, B. Noel, J.-M. Aury, Tara Oceans Coordinators, C. de Vargas, C. Bowler, E. Karsenti, E. Pelletier, P. Wincker, O. Jaillon, Functional repertoire convergence of distantly related eukaryotic plankton lineages abundant in the sunlit ocean. Cell Genom 2, 100123 (2022).

50. J. M. Gasol, C. Cardelús, X. A. G. Morán, V. Balagué, I. Forn, C. Marrasé, R. Massana, C. Pedrós-Alió, M. M. Sala, R. Simó, D. Vaqué, M. Estrada, Seasonal patterns in phytoplankton photosynthetic parameters and primary production at a coastal NW Mediterranean site. Scientia Marina 80, 63–77 (2016).

51. S. J. Bograd, M. P. Buil, E. D. Lorenzo, C. G. Castro, I. D. Schroeder, R. Goericke, C. R. Anderson, C. Benitez-Nelson, F. A. Whitney, Changes in source waters to the Southern California Bight. Deep Sea Research Part II: Topical Studies in Oceanography 112, 42–52 (2015).

52. Z. Yang, J. P. Bielawski, Statistical methods for detecting molecular adaptation. Trends Ecol Evol 15, 496–503 (2000).

53. N. C. P. Center, NOAA’s Climate Prediction Center. https://origin.cpc.ncep.noaa.gov/products/analysis_monitoring/ensostuff/ONI_v5.php.

54. S. J. Giovannoni, H. J. Tripp, S. Givan, M. Podar, K. L. Vergin, D. Baptista, L. Bibbs, J. Eads, T. H. Richardson, M. Noordewier, M. S. Rappé, J. M. Short, J. C. Carrington, E. J. Mathur, Genome Streamlining in a Cosmopolitan Oceanic Bacterium. Science 309, 1242–1245 (2005).

55. F. M. Lauro, D. McDougald, T. Thomas, T. J. Williams, S. Egan, S. Rice, M. Z. DeMaere, L. Ting, H. Ertan, J. Johnson, S. Ferriera, A. Lapidus, I. Anderson, N. Kyrpides, A. C. Munk, C. Detter, C. S. Han, M. V. Brown, F. T. Robb, S. Kjelleberg, R. Cavicchioli, The genomic basis of trophic strategy in marine bacteria. Proceedings of the National Academy of Sciences 106, 15527–15533 (2009).

56. A. Dufresne, L. Garczarek, F. Partensky, Accelerated evolution associated with genome reduction in a free-living prokaryote. Genome Biol 6, R14 (2005).

57. N. G. Walworth, E. J. Zakem, J. P. Dunne, S. Collins, N. M. Levine, Microbial evolutionary strategies in a dynamic ocean. Proceedings of the National Academy of Sciences 117, 5943– 5948 (2020).

58. A. B. Chase, C. Weihe, J. B. H. Martiny, Adaptive differentiation and rapid evolution of a soil bacterium along a climate gradient. Proceedings of the National Academy of Sciences 118, e2101254118 (2021).

59. A. A. Larkin, A. C. Martiny, Microdiversity shapes the traits, niche space, and biogeography of microbial taxa. Environmental Microbiology Reports 9, 55–70 (2017).

60. J. A. Sohm, N. A. Ahlgren, Z. J. Thomson, C. Williams, J. W. Moffett, M. A. Saito, E. A. Webb, G. Rocap, Co-occurring Synechococcus ecotypes occupy four major oceanic regimes defined by temperature, macronutrients and iron. ISME J 10, 333–345 (2016).

61. C. Sriaporn, K. A. Campbell, M. J. Van Kranendonk, K. M. Handley, Bacterial and archaeal community distributions and cosmopolitanism across physicochemically diverse hot springs. ISME COMMUN. 3, 80 (2023).

62. G. Rocap, F. W. Larimer, J. Lamerdin, S. Malfatti, P. Chain, N. A. Ahlgren, A. Arellano, M. Coleman, L. Hauser, W. R. Hess, Z. I. Johnson, M. Land, D. Lindell, A. F. Post, W. Regala, M. Shah, S. L. Shaw, C. Steglich, M. B. Sullivan, C. S. Ting, A. Tolonen, E. A. Webb, E. R. Zinser, S. W. Chisholm, Genome divergence in two Prochlorococcus ecotypes reflects oceanic niche differentiation. Nature 424, 1042–1047 (2003).

63. C. S. Ward, C.-M. Yung, K. M. Davis, S. K. Blinebry, T. C. Williams, Z. I. Johnson, D. E. Hunt, Annual community patterns are driven by seasonal switching between closely related marine bacteria. ISME J 11, 1412–1422 (2017).

64. C.-E. T. Chow, R. Sachdeva, J. A. Cram, J. A. Steele, D. M. Needham, A. Patel, A. E. Parada, J. A. Fuhrman, Temporal variability and coherence of euphotic zone bacterial communities over a decade in the Southern California Bight. ISME J 7, 2259–2273 (2013).

65. H. Teeling, B. M. Fuchs, C. M. Bennke, K. Krüger, M. Chafee, L. Kappelmann, G. Reintjes, J. Waldmann, C. Quast, F. O. Glöckner, J. Lucas, A. Wichels, G. Gerdts, K. H. Wiltshire, R. I. Amann, Recurring patterns in bacterioplankton dynamics during coastal spring algae blooms. eLife 5, e11888 (2016).

66. M. L. Bendall, S. L. Stevens, L.-K. Chan, S. Malfatti, P. Schwientek, J. Tremblay, W. Schackwitz, J. Martin, A. Pati, B. Bushnell, J. Froula, D. Kang, S. G. Tringe, S. Bertilsson, M. A. Moran, A. Shade, R. J. Newton, K. D. McMahon, R. R. Malmstrom, Genome-wide selective sweeps and gene-specific sweeps in natural bacterial populations. ISME J 10, 1589–1601 (2016).

67. Y. Hwang, P. R. Girguis, Differentiated Evolutionary Strategies of Genetic Diversification in Atlantic and Pacific Thaumarchaeal Populations. mSystems 7, e01477–21.

68. J. W. Chandler, Y. Lin, P. J. Gainer, A. F. Post, Z. I. Johnson, E. R. Zinser, Variable but persistent coexistence of Prochlorococcus ecotypes along temperature gradients in the ocean’s surface mixed layer. Environ Microbiol Rep 8, 272–284 (2016).

69. M. Chafee, A. Fernàndez-Guerra, P. L. Buttigieg, G. Gerdts, A. M. Eren, H. Teeling, R. I. Amann, Recurrent patterns of microdiversity in a temperate coastal marine environment. The ISME Journal 12, 237–252 (2018).

70. Y. Okazaki, S.-I. Nakano, A. Toyoda, H. Tamaki, Long-Read-Resolved, Ecosystem-Wide Exploration of Nucleotide and Structural Microdiversity of Lake Bacterioplankton Genomes. mSystems 7, e0043322 (2022).

71. A. Meziti, D. Tsementzi, L. M. Rodriguez-R, J. K. Hatt, H. Karayanni, K. A. Kormas, K. T. Konstantinidis, Quantifying the changes in genetic diversity within sequence-discrete bacterial populations across a spatial and temporal riverine gradient. ISME J 13, 767–779 (2019).

72. W. Yan, S. Wei, Q. Wang, X. Xiao, Q. Zeng, N. Jiao, R. Zhang, Genome Rearrangement Shapes Prochlorococcus Ecological Adaptation. Appl Environ Microbiol 84, e01178–18 (2018).

73. G. C. Kettler, A. C. Martiny, K. Huang, J. Zucker, M. L. Coleman, S. Rodrigue, F. Chen, A. Lapidus, S. Ferriera, J. Johnson, C. Steglich, G. M. Church, P. Richardson, S. W. Chisholm, Patterns and Implications of Gene Gain and Loss in the Evolution of Prochlorococcus. PLOS Genetics 3, e231 (2007).

74. T. O. Delmont, A. M. Eren, Linking pangenomes and metagenomes: the Prochlorococcus metapangenome. PeerJ 6, e4320 (2018).

75. M. L. Coleman, M. B. Sullivan, A. C. Martiny, C. Steglich, K. Barry, E. F. Delong, S. W. Chisholm, Genomic islands and the ecology and evolution of Prochlorococcus. Science 311, 1768–1770 (2006).

76. S. J. Biller, P. M. Berube, D. Lindell, S. W. Chisholm, Prochlorococcus: the structure and function of collective diversity. Nat Rev Microbiol 13, 13–27 (2015).

77. M. Schauer, V. Balagué, C. Pedrós-Alió, R. Massana, Seasonal changes in the taxonomic composition of bacterioplankton in a coastal oligotrophic system. Aquat. Microb. Ecol. 31, 163– 174 (2003).

78. K. H. Boström, K. Simu, Å. Hagström, L. Riemann, Optimization of DNA extraction for quantitative marine bacterioplankton community analysis. Limnology and Oceanography: Methods 2, 365–373 (2004).

79. C. Rinke, J. Lee, N. Nath, D. Goudeau, B. Thompson, N. Poulton, E. Dmitrieff, R. Malmstrom, R. Stepanauskas, T. Woyke, Obtaining genomes from uncultivated environmental microorganisms using FACS–based single-cell genomics. Nat Protoc 9, 1038–1048 (2014).

80. B. Bushnell, bbushnell/BBTools, (2026); https://github.com/bbushnell/BBTools.

81. M. Martin, Cutadapt removes adapter sequences from high-throughput sequencing reads. EMBnet.journal 17, 10–12 (2011).

82. G. Benoit, P. Peterlongo, M. Mariadassou, E. Drezen, S. Schbath, D. Lavenier, C. Lemaitre, Multiple comparative metagenomics using multiset k-mer counting. PeerJ Comput. Sci. 2, e94 (2016).

83. R. Team, R: A language and environment for statistical computing. MSOR connections (2014).

84. D. Li, C.-M. Liu, R. Luo, K. Sadakane, T.-W. Lam, MEGAHIT: an ultra-fast single-node solution for large and complex metagenomics assembly via succinct de Bruijn graph. Bioinformatics 31, 1674–1676 (2015).

85. H. Li, R. Durbin, Fast and accurate short read alignment with Burrows-Wheeler transform. Bioinformatics 25, 1754–1760 (2009).

86. H. Li, B. Handsaker, A. Wysoker, T. Fennell, J. Ruan, N. Homer, G. Marth, G. Abecasis, R. Durbin, The Sequence Alignment/Map format and SAMtools. Bioinformatics 25, 2078–2079 (2009).

87. D. D. Kang, J. Froula, R. Egan, Z. Wang, MetaBAT, an efficient tool for accurately reconstructing single genomes from complex microbial communities. PeerJ 3, e1165 (2015).

88. J. Alneberg, B. S. Bjarnason, I. de Bruijn, M. Schirmer, J. Quick, U. Z. Ijaz, L. Lahti, N. J. Loman, A. F. Andersson, C. Quince, Binning metagenomic contigs by coverage and composition. Nat Methods 11, 1144–1146 (2014).

89. Y.-W. Wu, Y.-H. Tang, S. G. Tringe, B. A. Simmons, S. W. Singer, MaxBin: an automated binning method to recover individual genomes from metagenomes using an expectation-maximization algorithm. Microbiome 2, 26 (2014).

90. G. V. Uritskiy, J. DiRuggiero, J. Taylor, MetaWRAP-a flexible pipeline for genome-resolved metagenomic data analysis. Microbiome 6, 158 (2018).

91. P.-A. Chaumeil, A. J. Mussig, P. Hugenholtz, D. H. Parks, GTDB-Tk: a toolkit to classify genomes with the Genome Taxonomy Database. Bioinformatics 36, 1925–1927 (2019).

92. M. R. Olm, C. T. Brown, B. Brooks, J. F. Banfield, dRep: a tool for fast and accurate genomic comparisons that enables improved genome recovery from metagenomes through de-replication. ISME J 11, 2864–2868 (2017).

93. X. Feng, H. Cheng, D. Portik, H. Li, Metagenome assembly of high-fidelity long reads with hifiasm-meta. Nat Methods 19, 671–674 (2022).

94. D. D. Kang, F. Li, E. Kirton, A. Thomas, R. Egan, H. An, Z. Wang, MetaBAT 2: an adaptive binning algorithm for robust and efficient genome reconstruction from metagenome assemblies. PeerJ 7, e7359 (2019).

95. A. Chklovski, D. H. Parks, B. J. Woodcroft, G. W. Tyson, CheckM2: a rapid, scalable and accurate tool for assessing microbial genome quality using machine learning. Nat Methods 20, 1203– 1212 (2023).

96. D. H. Parks, C. Rinke, M. Chuvochina, P.-A. Chaumeil, B. J. Woodcroft, P. N. Evans, P. Hugenholtz, G. W. Tyson, Recovery of nearly 8,000 metagenome-assembled genomes substantially expands the tree of life. Nat Microbiol 2, 1533–1542 (2017).

97. J. M. Kelliher, Y. Xu, M. C. Flynn, M. Babinski, S. Canon, E. Cavanna, A. Clum, Y. E. Corilo, G. Fujimoto, C. Giberson, L. Y. D. Johnson, K. J. Li, P.-E. Li, V. Li, C.-C. Lo, W. Lynch, P. Piehowski, K. Prime, S. Purvine, F. Rodriguez, S. Roux, M. Shakya, M. Smith, S. Sarrafan, S. Cholia, L. A. McCue, C. Mungall, B. Hu, E. A. Eloe-Fadrosh, P. S. G. Chain, Standardized and accessible multi-omics bioinformatics workflows through the NMDC EDGE resource. Comput Struct Biotechnol J 23, 3575–3583 (2024).

98. BBMap, SourceForge (2025). https://sourceforge.net/projects/bbmap/.

99. S. Nurk, D. Meleshko, A. Korobeynikov, P. A. Pevzner, metaSPAdes: a new versatile metagenomic assembler. Genome Res 27, 824–834 (2017).

100. D. H. Parks, M. Imelfort, C. T. Skennerton, P. Hugenholtz, G. W. Tyson, CheckM: assessing the quality of microbial genomes recovered from isolates, single cells, and metagenomes. Genome Res 25, 1043–1055 (2015).

101. A. Bankevich, S. Nurk, D. Antipov, A. A. Gurevich, M. Dvorkin, A. S. Kulikov, V. M. Lesin, S. I. Nikolenko, S. Pham, A. D. Prjibelski, A. V. Pyshkin, A. V. Sirotkin, N. Vyahhi, G. Tesler, M. A. Alekseyev, P. A. Pevzner, SPAdes: a new genome assembly algorithm and its applications to single-cell sequencing. J Comput Biol 19, 455–477 (2012).

102. M. Kolmogorov, D. M. Bickhart, B. Behsaz, A. Gurevich, M. Rayko, S. B. Shin, K. Kuhn, J. Yuan, E. Polevikov, T. P. L. Smith, P. A. Pevzner, metaFlye: scalable long-read metagenome assembly using repeat graphs. Nat Methods 17, 1103–1110 (2020).

103. R. Vaser, I. Sović, N. Nagarajan, M. Šikić, Fast and accurate de novo genome assembly from long uncorrected reads. Genome Res. 27, 737–746 (2017).

104. H. Li, Minimap2: pairwise alignment for nucleotide sequences. Bioinformatics 34, 3094–3100 (2018).

105. P. Danecek, J. K. Bonfield, J. Liddle, J. Marshall, V. Ohan, M. O. Pollard, A. Whitwham, T. Keane, S. A. McCarthy, R. M. Davies, H. Li, Twelve years of SAMtools and BCFtools. GigaScience 10, giab008 (2021).

106. S. T. N. Aroney, R. J. P. Newell, J. N. Nissen, A. P. Camargo, G. W. Tyson, B. J. Woodcroft, CoverM: read alignment statistics for metagenomics. Bioinformatics 41 (2025).

107. E. Garrison, G. Marth, Haplotype-based variant detection from short-read sequencing. arXiv arXiv:1207.3907 [Preprint] (2012). 10.48550/arXiv.1207.3907.

108. C. Sjöqvist, L. F. Delgado, J. Alneberg, A. F. Andersson, Ecologically coherent population structure of uncultivated bacterioplankton. ISME J 15, 3034–3049 (2021).

109. J. Shaw, J.-S. Gounot, H. Chen, N. Nagarajan, Y. W. Yu, Floria: fast and accurate strain haplotyping in metagenomes. Bioinformatics 40, i30–i38 (2024).

110. M. Borry, maxibor/floria-strainer, (2024); https://github.com/maxibor/floria-strainer.

111. T. Seemann, Prokka: rapid prokaryotic genome annotation. Bioinformatics 30, 2068–2069 (2014).

112. Y. Shimoyama, pyGenomeViz: A genome visualization python package for comparative genomics, (2024); https://github.com/moshi4/pyGenomeViz.

113. PPanGGOLiN: Depicting microbial diversity via a partitioned pangenome graph | PLOS Computational Biology. https://journals.plos.org/ploscompbiol/article?id=10.1371/journal.pcbi.1007732.

114. M. Pilgrim, Dive Into Python *3* (CreateSpace, 2010).

115. B. J. Knaus, N. J. Grünwald, vcfr: a package to manipulate and visualize variant call format data in R. Mol Ecol Resour 17, 44–53 (2017).

116. J. Oksanen, vegan: Community Ecology Package. R package version 2.0-10, edn. (2014).

