## Supplementary figures for "Decadal signatures of seasonal and ENSO-driven selection in microbial populations in distant oceans"

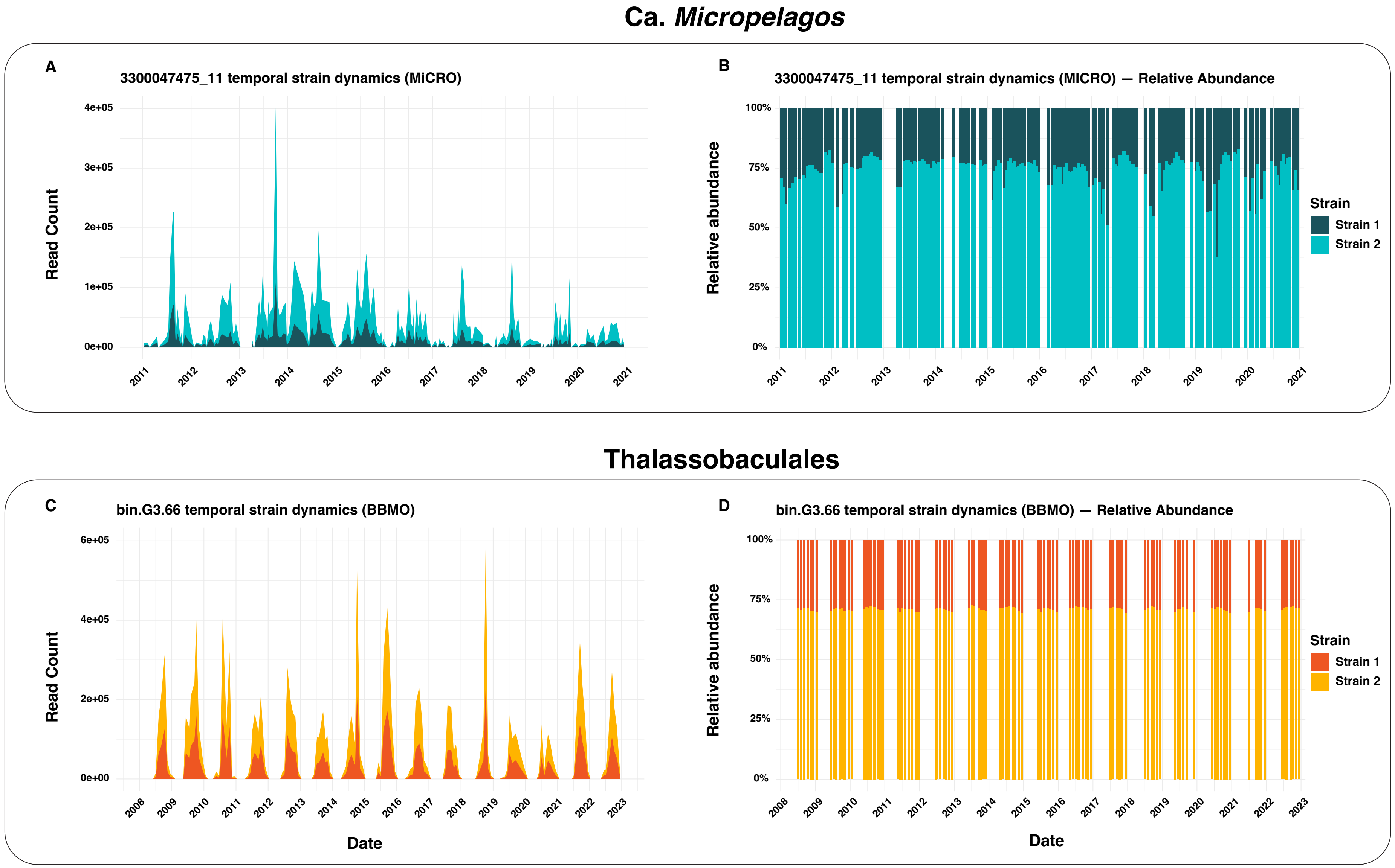


**Figure S1.** Strain-level dynamics of representative shared genomes with strain detection at only one site. **Panels A and B.** Temporal strain dynamics of a Ca. *Micropelagos* genome in MiCRO, shown as normalized read counts (A) and relative abundance (B). **Panels C and D.** Temporal strain dynamics of a Thalassobaculales genome in BBMO, shown as normalized read counts (C) and relative abundance (D). MiCRO genomes are shown in blue and BBMO genomes in orange; Strain 1 is represented by the darker shade of each color, and Strain 2 by the lighter shade. These examples complement Figure 5 by illustrating stable multi-strain dynamics in genomes where strain resolution was detected in only one of the two microbiomes.

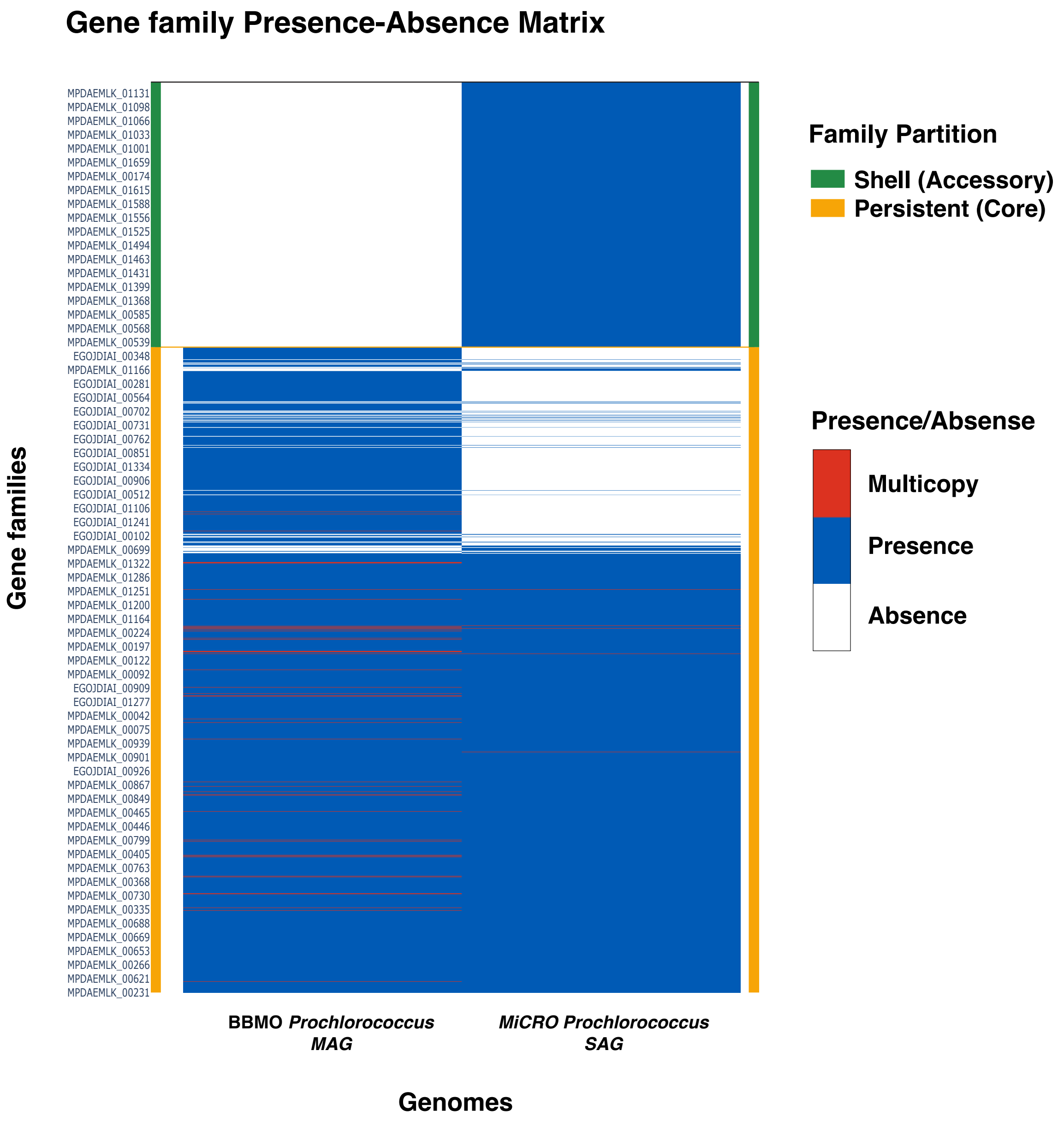


**Figure S2. Gene family presence-absence matrix for the shared *Prochlorococcus* genomes from BBMO and MiCRO.** Rows represent gene families and columns represent the BBMO long-read MAG and the MiCRO SAG. Rows are labeled using representative gene locus tags assigned by PPanGGOLiN, derived from the input genome annotations. Gene families are colored according to PPanGGOLiN partitions: persistent (core genes broadly conserved across the species, yellow) and shell (accessory genes present in a subset of genomes, green). Presence-absence of each family in a genome is indicated by color: blue = present (single copy), white = absent, and red = multicopy. Shared genes fall almost entirely within the persistent partition, while additional gene families were uniquely detected in one genome. The BBMO genome contains a higher number of multicopy families than MiCRO genome. These patterns highlight a highly conserved core genome alongside site-specific gene content variation.

### Flavobacteriaceae variant analysis

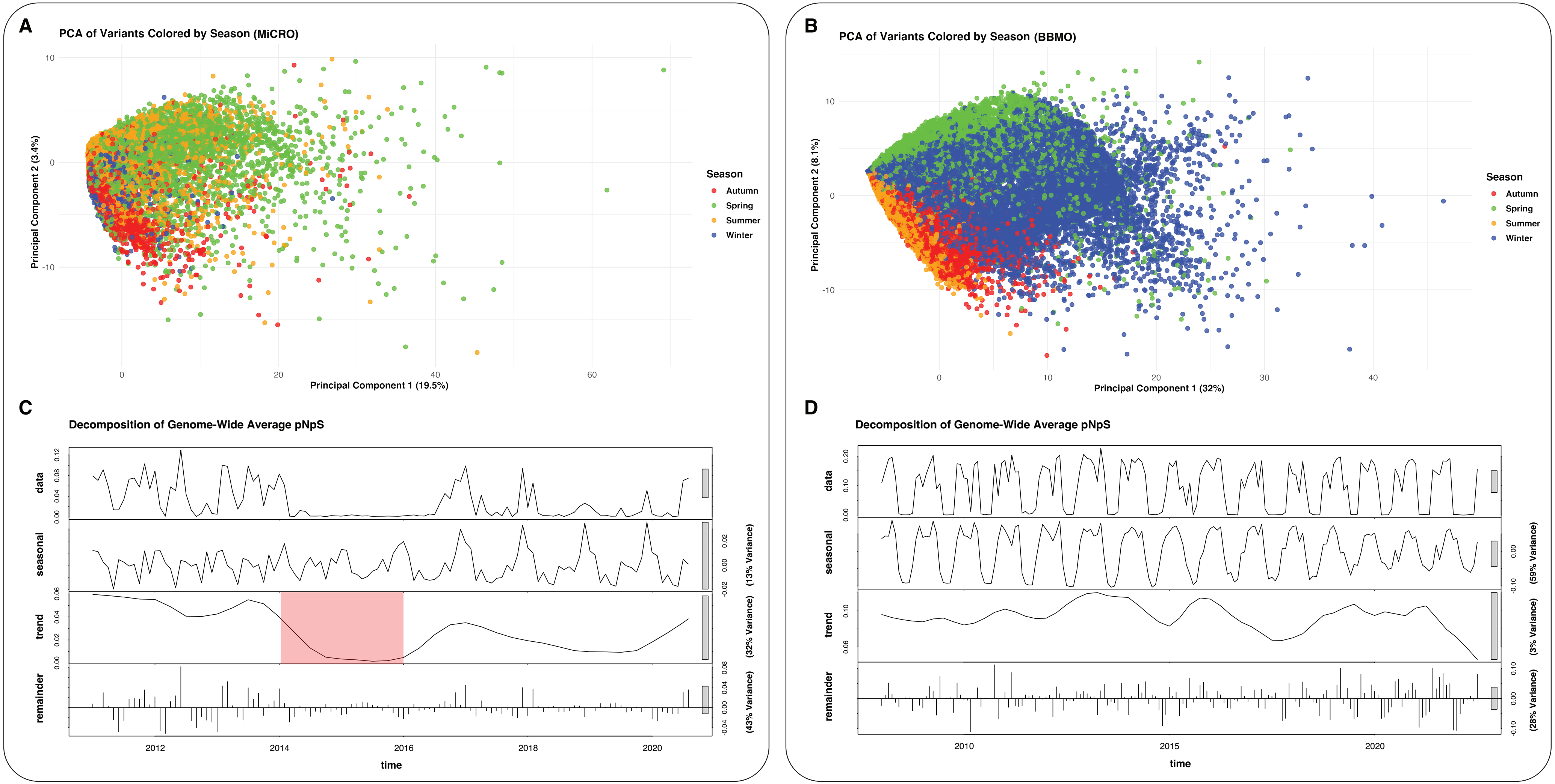

**Figure S3. Seasonal and long-term population dynamics of shared Flavobacteriaceae genomes at BBMO and MiCRO. Panels A and B.** Principal Component Analysis (PCA) of variants detected in the Flavobacteriaceae genomes from MiCRO (A) and BBMO (B). Each point represents a genetic variant or mutation detected in the corresponding VCFs after the variant calling analysis, colored according to the season where they appear the most (spring = green, summer = orange, autumn = red, winter = blue). Seasonal structure is visible, especially along PC1, indicating cyclic shifts in genetic composition. **Panels C and D.** Seasonal-Trend decomposition (STL) of the genome-wide average pN/pS ratio through time for the same Flavobacteriaceae genomes in MiCRO (C) and BBMO (D). For each site, the top panel ("data") shows the observed genome-wide average pN/pS ratio in each time series, followed by the seasonal, trend, and remainder components. The seasonal panels capture annual cycles in selection and population replacement; the trend component in MiCRO shows lower pN/pS values during El Niño years, while in BBMO the trend is minimal. The proportion of variance explained by seasonal and trend components is indicated next to each panel. In contrast to *Prochlorococcus* (Figure 7), Flavobacteriaceae showed the opposite long-term association with ENSO, highlighting that large-scale climatic variability can affect marine microbes in different ways at the population-genomic level.
